# Gene conversion drives allelic dimorphism in two paralogous surface antigens of the malaria parasite *P. falciparum*

**DOI:** 10.1101/2023.02.27.530215

**Authors:** Brice Letcher, Sorina Maciuca, Zamin Iqbal

## Abstract

While the malaria parasite *P. falciparum* has low average genome-wide diversity levels, likely due to its recent introduction from a gorilla-infecting ancestor (∼10,000-50,000 years ago), some genes display extremely high diversity levels. In particular, certain proteins expressed on the surface of human red-blood-cell-infecting merozoites (merozoite surface proteins, MSPs) possess exactly two deeply-diverged allelic forms that have not recombined. This phenomenon, called allelic dimorphism, is of considerable interest, but its origin and maintenance remains unknown.

In this study, we analysed the dimorphism in two highly-variable and paralogous MSPs, DBLMSP and DBLMSP2. Despite thousands of available Illumina WGS datasets from malaria-endemic countries, diversity in these genes has been hard to fully characterise as reads containing highly-diverged alleles fail to align to the reference genome. To solve this, we developed a pipeline leveraging genome graphs, enabling us to genotype them at high accuracy and completeness in comparison to a state-of-the-art GATK-based pipeline.

Using our newly-resolved sequences we found both genes are dimorphic in a specific protein domain (DBL), and that one of the two forms is shared across the genes. We identified clear evidence of non-allelic gene conversion between the two genes as the likely mechanism behind sharing, leading us to propose a new model for allelic dimorphism through gene conversion between diverged paralogs. This model is consistent with high diversity levels in these two genes despite the strong historical *P. falciparum* transmission bottleneck.

## Introduction

*Plasmodium falciparum* (hereafter, *Pf*) is a single-celled eukaryotic parasite causing malaria disease in humans. Malaria burden remains high worldwide, with 241 million cases and 627,000 deaths in 2020 according to the WHO (1). The high burden is in part due to *Pf*’s ability to evade the human immune system, mediated by two main mechanisms (2). Firstly, cell-surface-exposed antigens targeted by the immune system consist of functionally redundant gene families. For example merozoites, the parasite life stage infecting human red blood cells (RBC), use different members of the Rh and EBA families for invasion (2), and different members of the *var, rifin*, and *stevor* families enable infected RBCs to bind to the host microvasculature (3). Secondly, surface antigens are highly diverse at the sequence and immunological level. In the *var* family, diversity is mainly generated by frequent recombination and gene conversion (sequence copy-pasting) events, occurring both between orthologs during sexual reproduction, and paralogs on the same genome during asexual replication (4–7). Many merozoite surface proteins (MSPs) display deeply-diverged forms in populations, likely maintained by balancing selection for immune escape (8,9).

Deeply-diverged sequences in the MSPs were first identified by targeted sequencing of laboratory lines and a small number of field isolates, e.g. in MSP1, MSP2, MSP3 and MSP6 (10–13). Interestingly, in specific regions of each of these genes, exactly two classes of deeply-diverged alleles were found and recombination between the classes rarely observed, a phenomenon called *allelic dimorphism*. The reason why exactly two forms exist, and not more, have been debated but not resolved (14). In addition, the existence of multiple deep dimorphisms is at odds with the overall low levels of diversity in *Pf*, likely due to its very recent origin (10-50,000 years ago) in humans from a common ancestor with gorilla-infecting *P. praefalciparum* (15–17).

More recently, Illumina whole-genome sequencing (WGS) data for thousands of *Pf* field samples worldwide has become available in an effort to enable genomics-based epidemiology for malaria, called MalariaGEN (18). Variant calls released by the MalariaGEN consortium are obtained by aligning sequencing reads to the *Pf* 3D7 reference genome (19) and running GATK, a popular variant caller (20). An initial study using these calls confirmed that several MSPs are among the most highly-variable in the *Pf* genome and possibly subject to balancing selection (9), but fully characterising them has remained challenging.

In particular, for dimorphic MSPs the non-reference alleles are subject to highly biassed ascertainment due to *reference bias*: reads from highly-diverged alleles fail to align to the reference and thus alleles which are very different from the reference genome are under-detected. Miles *et al*. showed that some diverged alleles could be reconstructed using an assembly-based method called Cortex (21). However, Cortex still misses some of these alleles and is not sensitive to small variants like SNPs; furthermore the study only characterised crosses of 6 laboratory strains, so that a comprehensive view of variation in the MSPs is still missing.

Here, we analyse two MSPs, DBLMSP and DBLMSP2, in detail using the WGS data from MalariaGEN (22). Both genes are cell-surface exposed antigens recognised by the human immune system (23,24) and are among the most highly-variable genes in *Pf* (excluding the *var*s) (21). They are part of an eight-gene tandemly arrayed family of paralogs, as identified from sequence sharing: all eight genes possess an N-terminal signal sequence, six (including DBLMSP and DBLMSP2) possess a C-terminal SPAM domain, and DBLMSP and DBLMSP2 further uniquely possess a DBL domain (24) (summarised in supp. fig. 1). DBL domains mediate a number of important malarial host-pathogen interactions, including between erythrocyte binding antigen (EBA) gene products and RBCs during invasion (25,24), and between *var* gene products on infected RBCs and various human receptors, enabling sequestration (26). However their function in DBLMSP and DBLMSP2 is still largely unknown (27).

To comprehensively genotype variants in these genes, we developed a pipeline leveraging different approaches, including using genome graphs to remove 3D7-alignment reference bias (28). Applying the pipeline to >3,500 global *Pf* samples, we show this approach can recover significantly more variation than previously possible, enabling us to analyse these genes at a new level of detail. We confirm extreme levels of diversity in DBLMSP and DBLMSP2, and identify a deep dimorphism in each spanning the DBL domain, with one of the two forms being shared between the two genes. We find evidence that sequence sharing is due to gene conversion between the paralogs, leading us to propose a new model for the generation and maintenance of allelic dimorphism, also consistent with the recent gorilla-to-human transmission bottleneck.

For the remainder of this paper, we refer to DBLMSP and DBLMSP2 collectively as the *DBs*.

## Results

### 1. New genotyping pipeline outperforms MalariaGEN’s GATK-based pipeline

To analyse variation in the DBs, we used data from the 2021 MalariaGEN *Pf* data release consisting of >7,000 samples (22). Of these, we retained 3,589 samples passing MalariaGEN’s quality-controls and showing evidence of being clonal (see Methods), as multiple infections are common in *Pf* and can confound genotyping (18,29). These samples come from a total of 29 countries (supp. fig. 1.1). After read preprocessing and quality control (see Methods), all 3,589 samples were genotyped using a newly-developed pipeline.

In Fig. 1, we illustrate MalariaGEN’s existing pipeline (panel a) and our new pipeline (panel b). Both approaches first genotype samples individually and then re-genotype each sample at the union of all variants found (joint genotyping). MalariaGEN uses GATK for both steps (30), while in our new pipeline per-sample calls are obtained using a range of tools and joint genotyping is performed using our genome-graph-based genotyper, gramtools (28). We call MalariaGen’s pipeline ’GATK-based pipeline’ and our new pipeline ’gramtools-based pipeline’, and provide a rationale for each of our pipeline’s steps in the Methods.

**Figure 1.**
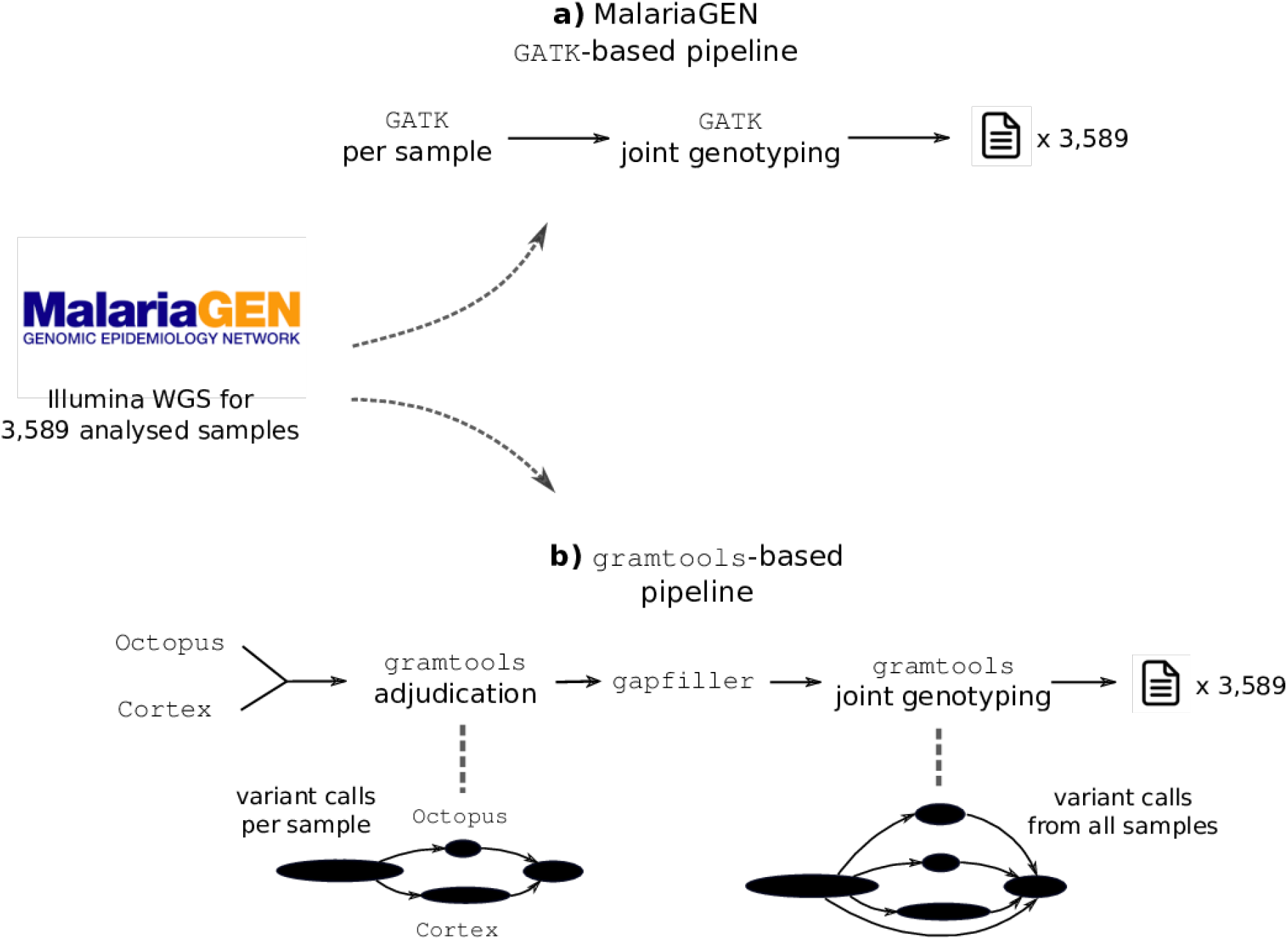
Existing and novel genotyping pipelines applied to the MalariaGEN data. Panel a illustrates MalariaGEN’s existing GATK-based pipeline, and panel b illustrates our new pipeline. Both first discover variants in each sample individually before re-genotyping each sample at the union of all variants. GATK relies on the linear reference genome to do this, while gramtools uses a genome graph.

To evaluate genotype calls, we designed an evaluation framework based on two different approaches: one using independent samples with both Illumina short reads and PacBio long reads, from which highly contiguous assemblies were built in (31), and another based on aligning reads back to the inferred genomes consisting of the reference genome with variant calls applied. We also used the latter approach to create a set of analysis-ready DBLMSP and DBLMSP2 sequences that we deemed ’confidently-resolved’, after applying stringent criteria for realigned read coverage, reference-agreement and insert size to the final output of each pipeline (see Methods). For the GATK-based pipeline, 49% (DBLMSP) and 12% (DBLMSP2) of all sequences were confidently-resolved, while for our new pipeline >81% were confidently-resolved for both genes (supp. fig. 1.2). Our new sequence set also contained much more variation, as we show in the next section.

### 2. The DBL domain of the DBs is highly variable and contains shared and private sequences

To analyse polymorphism levels in the DBs, we translated all high-quality gene sequences (i.e., passing all the filters above) into protein sequences, excluding sequences with one or more premature stop codons (234/6123 (3.8%) sequences for our pipeline; see methods). We then produced a multiple-sequence alignment (MSA) of the remaining DB protein sequences using mafft (32) and computed two measures of sequence diversity, shown in Fig. 2. In panel a, we show within-gene heterozygosity (y-axis), defined as the probability that, for a given gene and at a given aligned position (x-axis), two randomly chosen amino acids from the population differ. For both DBLMSP and DBLMSP2, much less diversity was recovered by the GATK-based pipeline (left-hand side panels) compared to ours (right-hand side). In our sequences only, a central region of each gene is particularly polymorphic and spans their DBL domain, delimited with blue vertical dotted lines (see methods for annotation).

**Figure 2.**
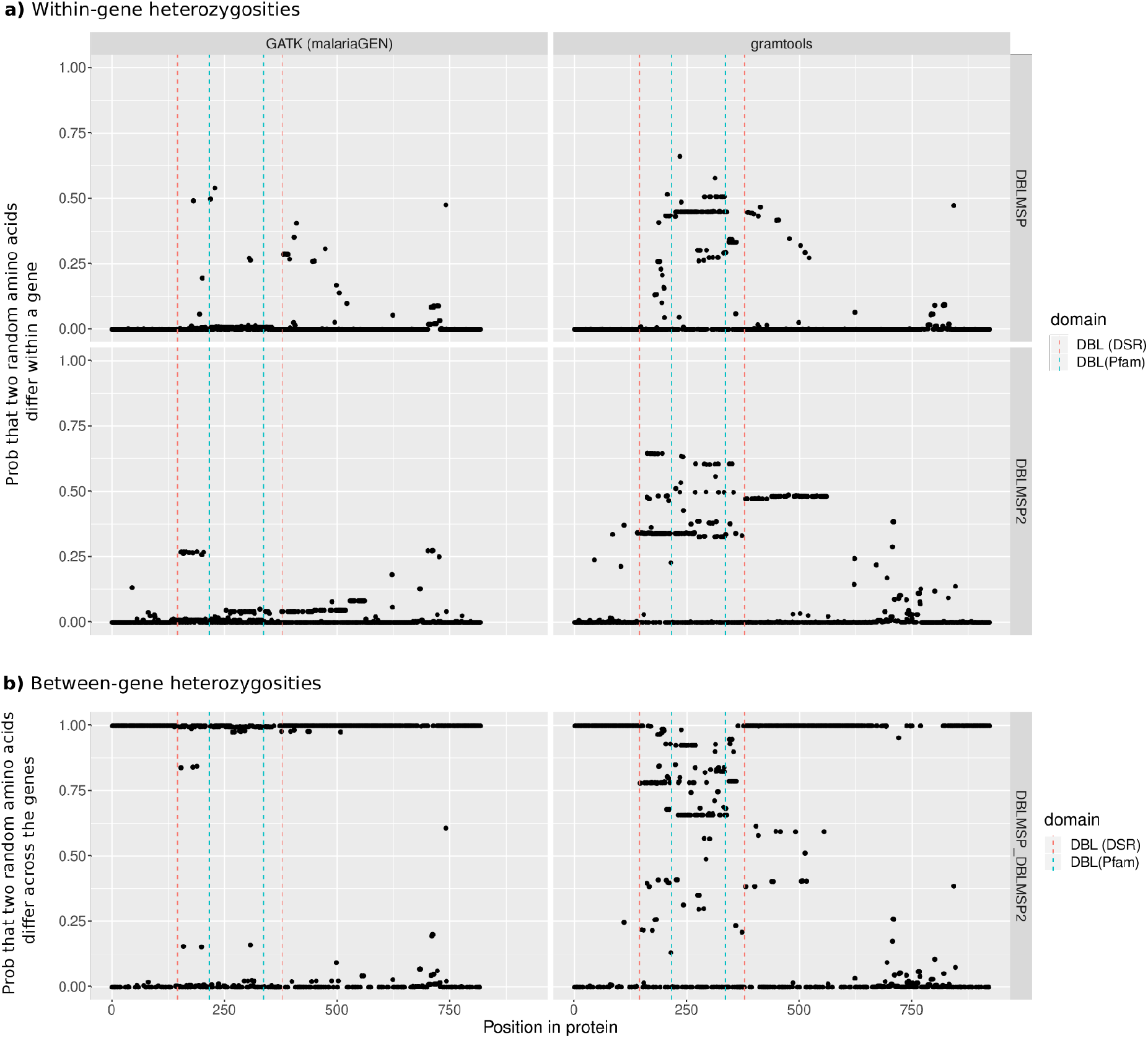
Within- and between-gene heterozygosities in DBLMSP and DBLMSP2. In panel a, the y-axis measures the probability that, at each aligned protein position (x-axis), two randomly-chosen amino acids differ, for each gene and each of the GATK- and gramtools-based pipelines. A region of extreme diversity spans the DBL domain, annotated with blue vertical dotted lines, and is only visible with our new pipeline. Panel b shows the probability that two randomly-chosen amino acids, one from each gene, differ. A value of 1 indicates no amino acids in common, i.e. full divergence of the two genes. The DBL domain lies in a region of shared sequence, where no amino-acid has fully diverged, and indicated with red vertical dotted lines - we call this the DBL-spanning region (DSR). We note that a smaller C-terminal region also displays positions with putative sequence sharing, but these are in fact gap characters in an indel-rich region of the alignment.

In panel b we show between-gene heterozygosity, defined as the probability that, for two genes at an aligned position, two randomly-chosen amino acids - one from each gene - differ. A value of 1 indicates no amino acids in common between the sequences of the two genes (fully-diverged position), while a value of 0 indicates a single identical amino acid is found in both genes (fixed position). While many positions are fully-diverged, a region spanning the DBL domain, shown with red vertical dotted lines, has no single fixed-differences between the genes, indicating sequence sharing. This observation is impossible with previous methods (i.e. the GATK-based pipeline). We call this region the DBL-spanning region, or DSR, and focus the rest of the analysis in this paper on this region, and using our gramtools-pipeline results.

To explore sharing patterns in the DSR in more detail, we broke each aligned sequence into peptides of length 10 (*10-mers*), and called a 10-mer *shared* if it was seen at least once in each gene at a given position in the MSA, and *private* if not. Shared peptides are abundant in populations, occurring at ∼25-50% frequencies along the entire DSR in all 16 countries with more than 50 samples (supp. fig. 2.1).

To visualise the shared and private sequences themselves, we separated the original MSA into three MSAs: one containing the private peptides of DBLMSP, one for those of DBLMSP2, and one for the shared peptides. We then built an HMM logo for each MSA, a visual representation in which amino acid frequency is encoded by letter height, shown in Fig. 3. Tracks labelled ’DBLMSP’ and ’DBLMSP2’ show the private peptides, and ’Both’ show the shared peptides. Either side of the DSR only private peptides are found, meaning the genes have diverged (as in Fig. 2). Inside the DSR, two amino acids are commonly found in each gene - one private and one shared - interspersed with fixed amino acids (e.g. the well-known ’PPRR’ motif of DBL domains, and a number of cysteine and tryptophan residues) and some more highly polymorphic locations (e.g. the C-terminal region of the DSR). Some low-frequency amino acids additionally exist, though not visible in Fig. 3 (see supplementary fig. 2.2).

**Figure 3.**
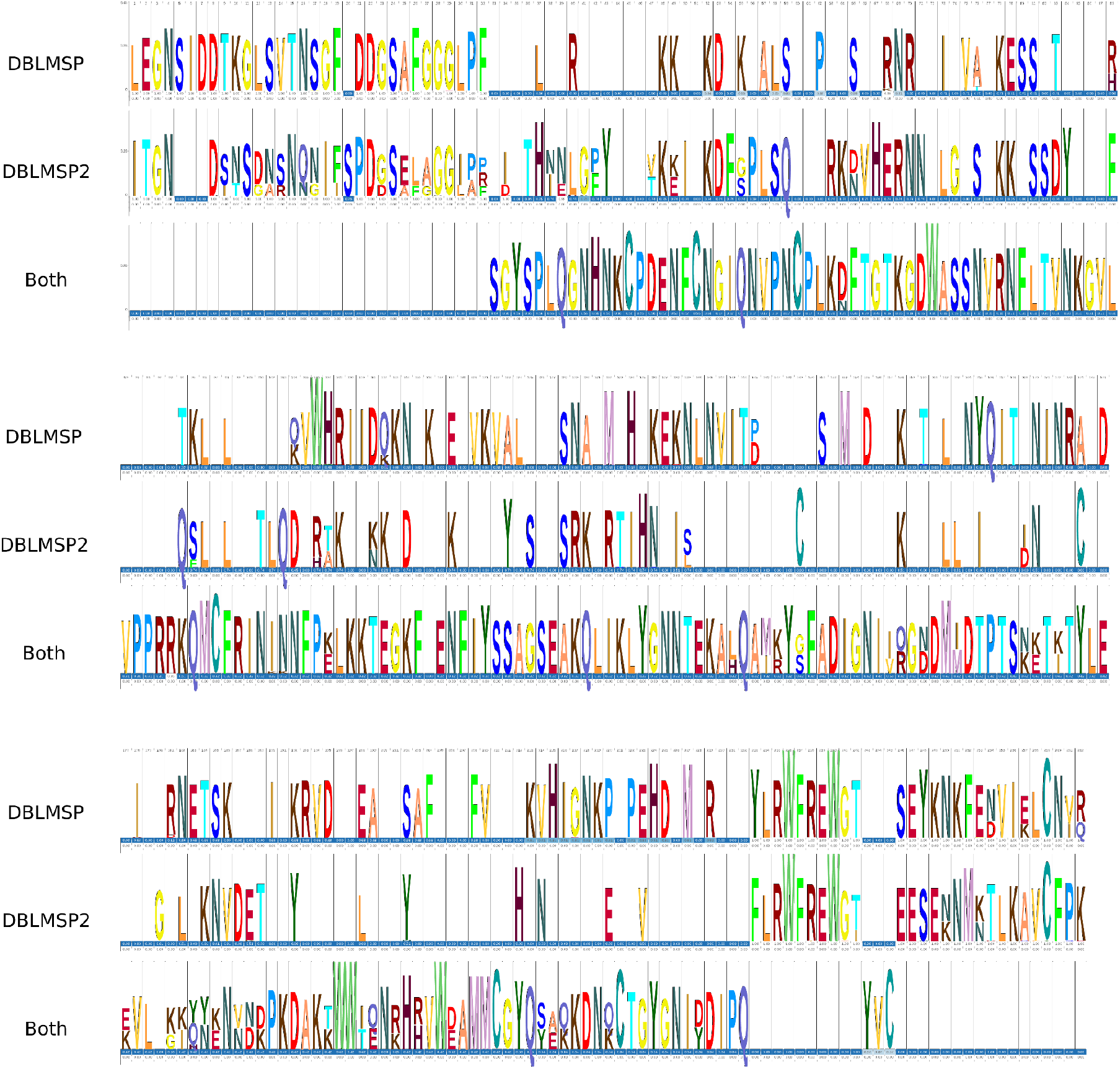
HMM logos of private and shared DB sequences in the DSR. One logo was produced for peptides found only in DBLMSP (top panel), only in DBLMSP2 (middle panel), and found on both genes (lower panel, labelled ’Both’). The three tracks are broken into segments for visual clarity. At each position, observed amino acids are shown, with letter height proportional to amino acid frequency. There is mostly one prototypical private form for each gene (first two tracks) and one prototypical shared form (or two, in the C-terminal half of the protein domain), indicating dimorphism in this region.

The motifs in Figure 3 suggest the DSR of each gene is dimorphic and that, in contrast to other dimorphic MSPs in *Pf*, here one of the two forms is shared between the genes. In the other dimorphic MSPs, recombination between the two forms has been reported to be rare or absent (13,33). In the next section, we map out recombination patterns in the DSR to also test this, and additionally identify what drives sequence sharing.

### 3. Recombination and gene conversion in the DBL domain

To detect recombination in our DB protein sequences, we used a method developed by Zilversmit *et al*. (34) for studying the *var* genes in *Pf*. Briefly, each sequence in a panel is aligned to all others using a HMM-based model that performs pairwise alignment between target and donor while allowing for switching between donors (i.e. recombination). Given our high number of sequences we first clustered them into 35 representatives (at 96% identity), as sequences that are too closely identical would only get aligned to each other, obscuring more distant recombination. We ran Zilversmit *et al*.’s implementation MosaicAligner on this panel, and built visual representations of the outputs to verify each inferred breakpoint (code available with this paper, see data availability). In Figure 4 panel a, we show one such ’mosaic alignment’ in detail. The target, a DBLMSP sequence (second row), is a recombinant of two other DBLMSP donors, and the vertical red line shows the inferred recombination breakpoint. Either side of the breakpoint, the highlighted donor has fewer mismatches (red-coloured letters) than the non-highlighted one; this held for all 35 alignments (supp. fig. 3.1). In panel b, we show three full mosaic alignments: one for the first representative of each gene, and one for which the breakpoints span donors on both genes (discussed next). Illustrations for all 35 mosaic alignments are available with this paper (see data availability). Overall, across the 35 alignments we found breakpoints at a total of 13 (DBLMSP) and 15 (DBLMSP2) of 254 positions in the DSR, with no apparent hotspot structure as they were spread across the full region (panel c).

**Figure 4.**
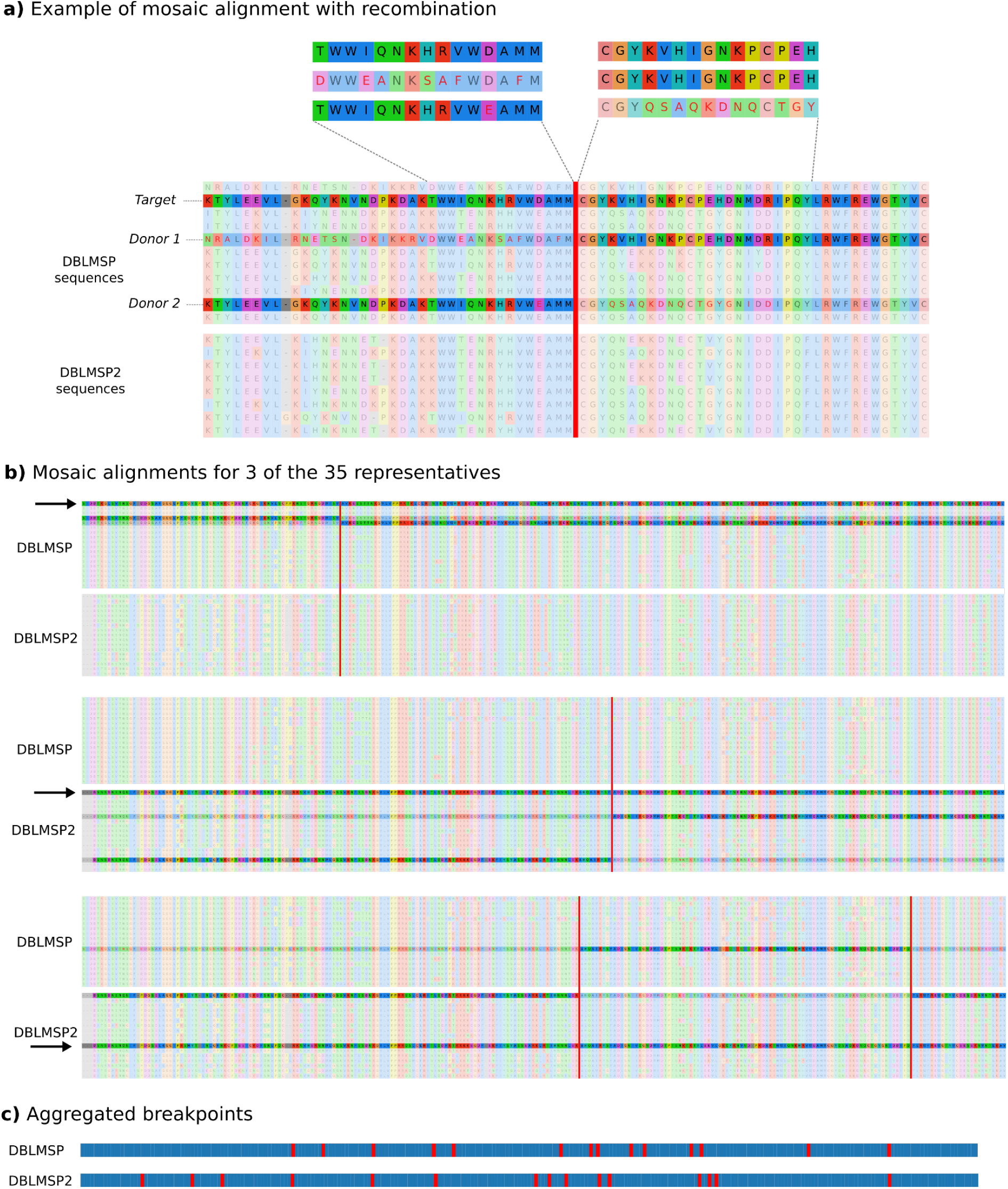
Mosaic alignments reveal widespread recombination in the DBs. a) Visual confirmation of MosaicAligner’s inferred alignment for the first DBLMSP sequence (*target*; fully-shaded sequence), aligned to two other DBLMSP sequences (*donors*; partially-shaded sequences). The red vertical line marks the switch in alignment between the two donors. Either side of the switch, the target aligns to the shaded donor with many fewer edits than to the other donor (red-coloured letters flag mismatches). b) full mosaic alignments illustrated for three out of 35 representative sequences in the panel. In each panel, the aligned target is the fully-opaque row labelled with an arrow, and the donors are shown as partially-opaque rows. Illustrations for all 35 alignments are available with this paper. c) For each gene, the aggregated locations of all breakpoints from the mosaic alignments are shown. The breakpoints did not seem to cluster into hotspots.

For three of the mosaic alignments, the donor sequences came from different genes (last alignment of Fig. 4 panel b and supp. fig. 3.2), consistent with sequence exchange between the paralogs. These can occur when repairing double-strand breaks, using another genome, e.g. during recombination in meiosis, or the same genome, e.g. during mitosis, and can lead to sequence pasting from the unbroken template, also called gene conversion (35). We tested for within-genome gene conversion between the DBs as a possible basis for sequence sharing in the DSR. For each of the 2,882 samples in which both DB genes were confidently-resolved, we pairwise aligned the two genes and measured sequence identity at the DNA codon level (see methods). In Figure 5, we illustrate the 209 samples for which the fraction of identical codons in the DSR was high (>0.5; see supp. fig. 3.3 for full distribution), after ruling out duplications of the DSR having occurred inside a gene (see methods). Each row shows one sample’s DB sequence alignment in the DSR and each column shows one codon, with cells coloured beige for identical codons and black for different codons. Stretches of near-uninterrupted beige are clearly visible, supporting the occurrence of within-genome gene conversion between the DBs, as opposed to between-genome conversion that would not result in identical sequences on the same genome.

**Figure 5.**
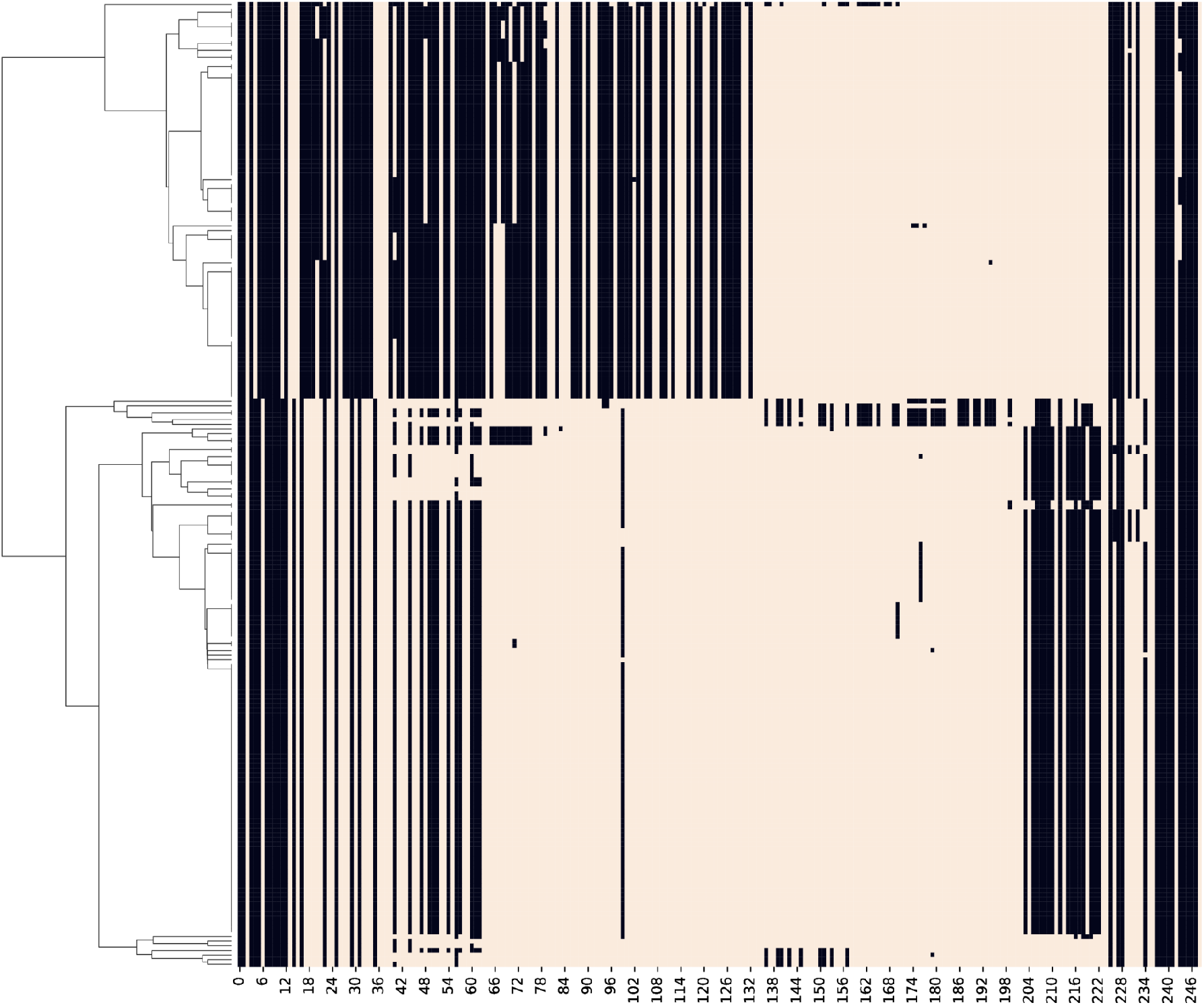
Evidence for within-genome, non-allelic gene conversion between DBLMSP and DBLMSP2 in the DSR. For each of the samples in which both DB gene sequences were confidently-resolved, we aligned their DNA sequences and recorded the fraction of identical codons in the DSR. Rows show the 209 samples with a value > 0.5, and columns show each codon position. Cells are coloured beige for identical codons and black for different codons. Strips of near-all beige indicate likely sequence copying between the two genes in a sample and support gene conversion occurring within each genome. Hierarchical clustering of the row values distinguishes two main clusters (left-hand tree), corresponding to different conversion breakpoints.

Clustering the identity matrix in Figure 5 (left-hand side tree) distinguishes two main conversion clusters; these are also different at the sequence level (supp. fig. 3.4). Both clusters were geographically widespread, occurring across both west and east sub-saharan Africa and South-East Asia (supp. fig. 3.5), suggesting they are being actively maintained, either through selection or recurrent conversion.

We now bring together the data from Figures 3, 4 and 5 into a single summary figure. In the discussion, we use these data to propose a model for the generation and maintenance of allelic dimorphism in the DBs.

### 4. Summary view of sequence relationships in the DBs

In Figure 6, we show a hierarchical clustering tree built from all unique protein sequences in the MSA of the DBs. The tree clearly shows three main clades, marked A, B and C. Clades A and C consist exclusively of DBLMSP (yellow in innermost coloured ring) and DBLMSP2 (blue) sequences respectively, while clade B contains sequences from both genes. The middle ring measures the fraction of shared (i.e. observed in both DBLMSP and DBLMSP2) 10-mer peptides of each sequence and clearly shows clade B is the ’shared form’, with all members having high shared sequence content (green). The recombination events illustrated in Fig. 4 are shown here as dotted lines joining sequence pairs inferred to have recombined. Almost all recombinations occur inside of the three main clades, although a few recombinations occur across clades as well (see supp. fig. 4.1 for one example). Finally, the outermost ring shows sequences belonging to the two gene conversion clusters in Fig. 5, with colours indicating the fraction of identical codons between the two DB sequences in that sample. The two conversion clusters are enriched in different sub-clades. Notably, conversion cluster 1 in Fig. 5, with about 50-60% identity, occurs inside two sub-clades, one inside sub-clade ’B.2’, and another in sub-clade ’A.1’.

**Figure 6.**
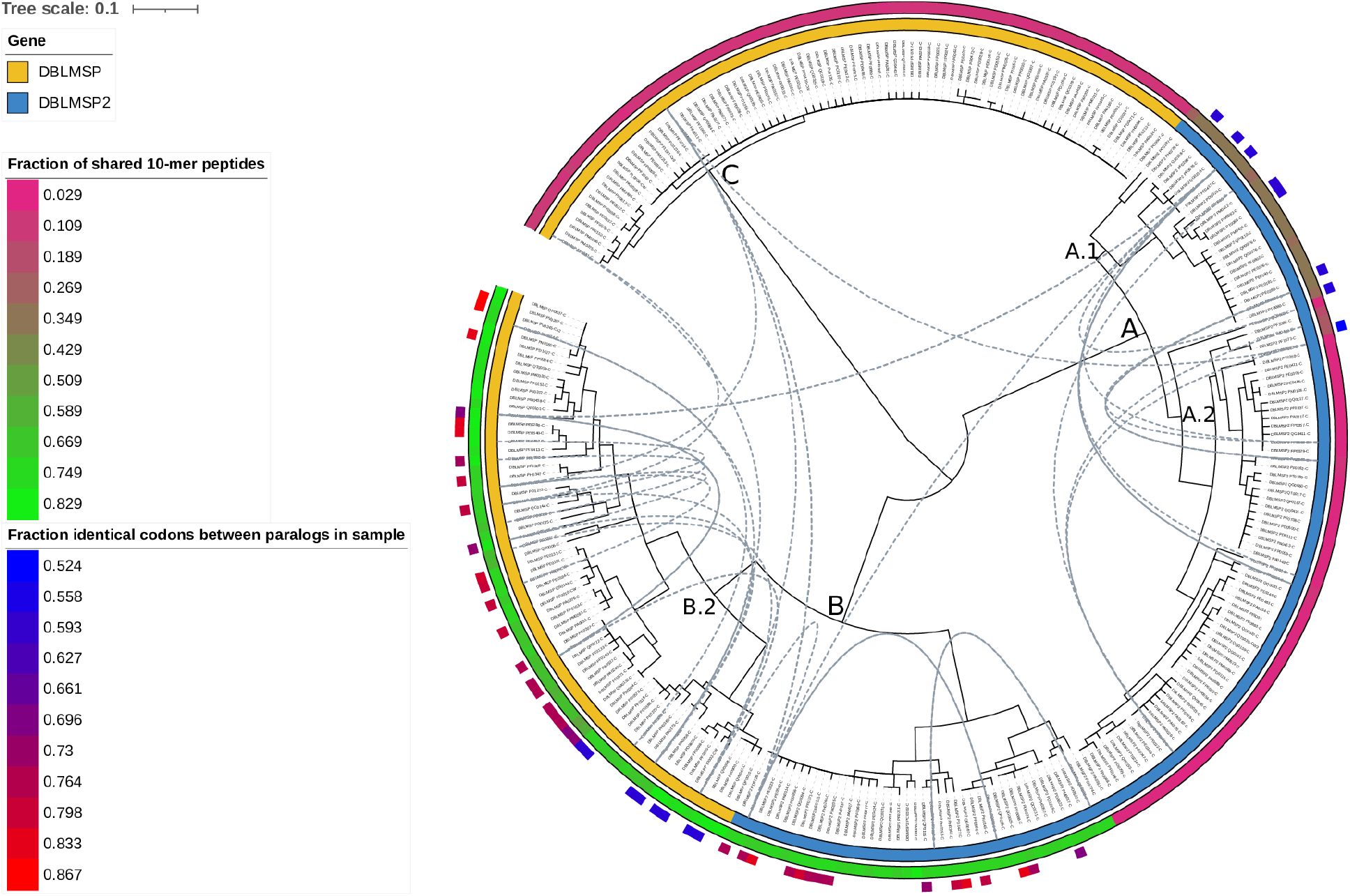
Allelic dimorphism, recombination and gene conversion in the DBs. A hierarchical clustering tree of all unique DBL-spanning protein sequences was built. The innermost ring colours sequences by gene of origin (DBLMSP, DBLMSP2). The next ring moving outwards colours sequences by the fraction of 10-mer peptides that are shared. The outermost ring shows samples in the conversion clusters shown in fig. 5, and colours sequences by the fraction of identical codons between DBLMSP and DBLMSP2 in a sample. Dotted lines link samples that recombined from the ’mosaic’ analysis (Fig. 4). Three main clades appear in the tree, marked A, B and C, and correspond to the private (A, C) and shared forms in Fig. 3. Recombination mostly occurs within each clade/form. A subform inside the DBLMSP2-private clade exists with higher sharing, and matches with gene conversion cluster 1 from Fig. 5. Subclades exist within the shared clade as well.

**Figure 7.**
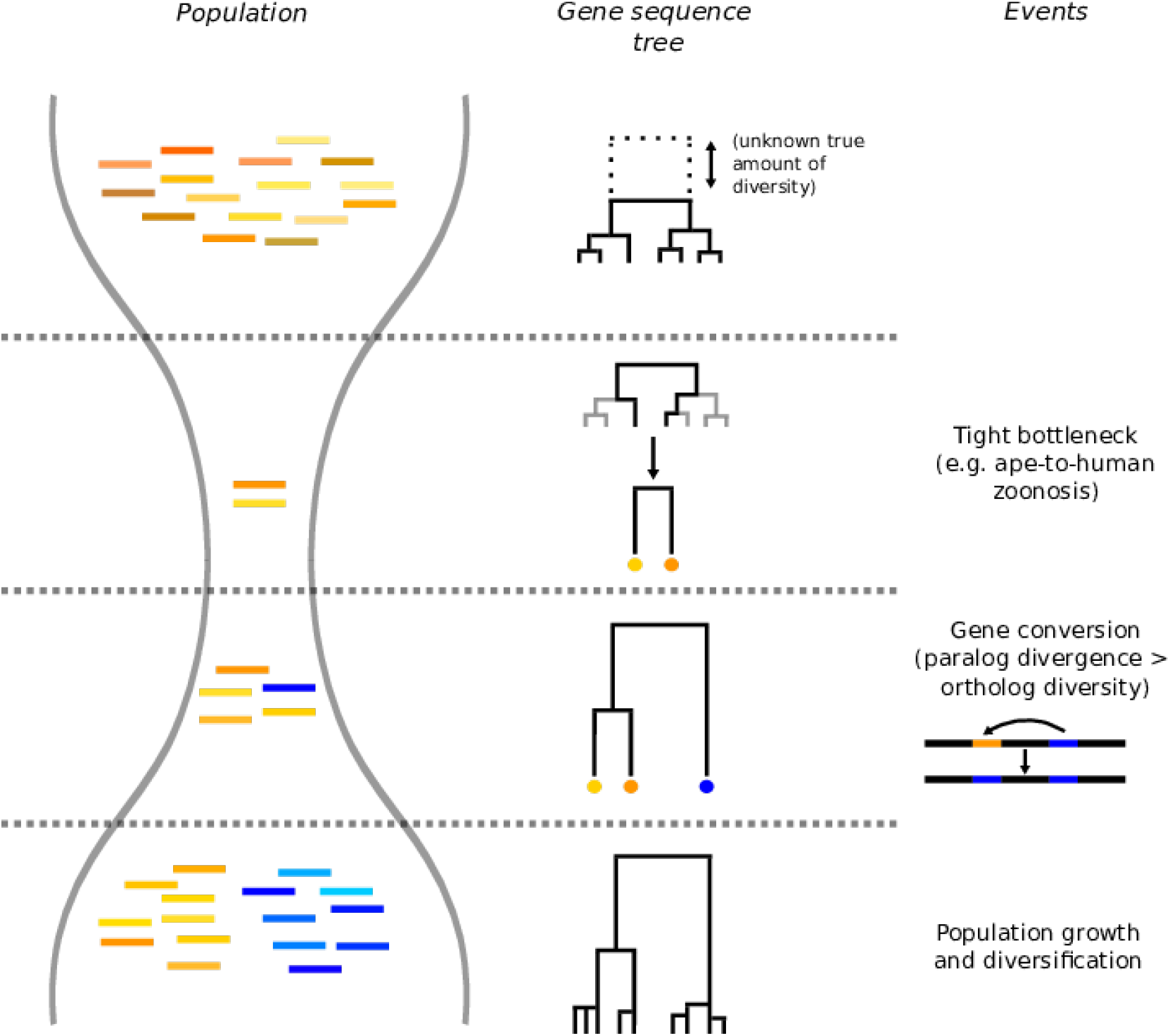
A model for allelic dimorphism through gene conversion between diverged paralogs. On the left, a population of gene sequences is shown, with the corresponding sequence tree to its right (one gene only). Time goes from top to bottom, and key events are depicted on the far-right. From an initial population with unknown levels of true diversity, a strong bottleneck occurs, consistent with a recent gorilla-to-human zoonosis from a single or a few individuals. We then propose dimorphism was introduced by gene conversion from a nearby paralog, as observed between DBLMSP and DBLMSP2. After population growth, diversification and balancing selection, exactly two highly-diverged, non-recombined lineages are observed in the present.

In the discussion, and after testing for direct evidence of gene conversion below, we propose a gene-conversion based model for allelic dimorphism consistent with our data.

### 5. Testing for direct evidence of evolution in the DBs

The recombination and gene conversion events studied so far were inferred indirectly from population-level data. To test for direct evidence of these events in the DBs, we also analysed data from repeatedly-sequenced isolates through time. We looked for mutations in the DBs in two sources: the ’clone trees’ from Hamilton et al. (36), who repeatedly cloned, cultured and sequenced individual isolates (spanning ∼700 erythrocytic life cycles in total), and four experimental genetic crosses between different strains of *Pf* (142 sequenced parents and progeny in total (21,37); see data availability and methods for details). Overall we found just two point mutations in one of the genetic cross progeny and no direct evidence of recombination or gene conversion in the DBs (supp. fig. 5.1). We note, though, that gene conversion in genes RH2a and RH2b was observed in a repeatedly cultured and sequenced isolate by Cortes (38).

## Discussion

The existence of allelic dimorphism in specific *P. falciparum* (*Pf*) genes is a long-standing puzzle. While highly-diverged forms in merozoite surface proteins (MSPs) may occur due to balancing selection via immune pressure, this does not predict that exactly two forms should be observed (and not three, for example). Further, while recombination within forms has been observed in MSPs (including here in the DBs) (33), recombination-induced intermediates between emerging forms are expected, but have not been found. In 2008, Roy *et al*. proposed a number of models to explain these observations but concluded that none were satisfactory and more data was needed (14).

A second puzzle for MSPs revolves around the recently-resolved origin of *Pf*. Current data strongly supports a recent introduction of *Pf* into humans, about 10-50,000 years ago, by zoonosis from a common ancestor with the gorilla-infecting *P. praefalciparum* (15). This is consistent with the very low genome-wide diversity levels observed in *Pf* (∼10-fold lower than other species of the sub-genus *Laverania* (17,39)), possibly through the jump of only one or a few individuals into human (17,40). However this bottleneck is at odds with the existence of dimorphic loci. The existing argument is that such highly diverged haplotypes must have existed prior to the split, and for both forms to survive the host jump, there must have either been a multiple-infection (not too small bottleneck) or multiple independent jumps (16).

Our work in this study contributes to both these debates, by revealing an alternate mechanism for generating dimorphism in the DBs. We genotyped the highly-variable and paralogous genes DBLMSP and DBLMSP2 (abbreviated DBs) using a newly-developed pipeline applied to Illumina data from >3,500 global *Pf* samples. The pipeline could resolve many more gene sequences compared to those released by the MalariaGEN consortium using GATK, enabling us to analyse segregating variation in >6,000 confidently-resolved DB sequences. Confirming previous reports, we found that both DBLMSP and DBLMSP2 are extremely polymorphic, especially in a central region spanning the DBL domain inside which many positions have protein-level heterozygosities >0.5. In common with other merozoite surface proteins (MSPs), we found that the DBs are dimorphic: two highly-diverged forms exist in each gene, and recombination between the forms is rare.

In Fig. 7, we illustrate a new model for explaining allelic dimorphism in the DBs. Looking at the sequence tree of a single gene (middle column), and starting with an ancestral population of unknown size, a small number of lineages are sampled at the point of gorilla-to-human zoonosis (second row; two sampled lineages in the example), leading to a highly reduced gene tree by bottleneck. We propose dimorphism can then be created in the tree by gene conversion from a paralog, leading to two highly-diverged lineages (third row). This can arise if the divergence between the paralogs exceeds the (orthologous) diversity levels within each gene. This is expected under neutral evolution, and we found it applied to the DBs in practice (supp. fig. D.1). Subsequent population growth, diversification and balancing selection would then lead to the current observed tree (fourth row). The absence of inter-form recombination could then be due to the low homology level between the two lineages and the short amount of time since zoonosis. An interesting parallel to our model is one of the Roy *et al*. models, in which orthologous genes diverged after speciation (e.g. between human and gorilla), and interspecific hybridisation introduced a diverged gene sequence into *Pf*. Our model provides a simpler, ’single-genome’ alternative, also consistent with only 3 clades in the joint allele tree of both paralogs, rather than 4 expected for interspecific hybridisation. A slightly more complicated scenario is required for explaining the full observed data in the DBs (Fig. 6; three lineages in total), which we provide in supp. fig. D.2.

In the future, this model could be tested in other dimorphic MSPs: notably, dimorphic gene MSP2 occurs in tandem with another MSP (MSP4), and dimorphic genes MSP3 and MSP6 both occur in the same eight-gene paralog tandem array as DBLMSP and DBLMSP2 (supp. fig. 1). Interestingly, and as for the DBs here, Nielsen *et al*. (2003) reported gene conversion between the paralogous genes FP2A and FP2B located 10kbp apart, causing the genes to look far more diverse than consistent with a recent bottleneck (41). We do not propose that this model explains all dimorphisms, however: other dimorphic genes like MSP1 or EBA-175 do not occur in tandem with a paralog (42,43). More generally, our model does not imply that exactly two forms should exist, nor that they do not recombine, both of which would be hard to expect in the long term.

Gene conversion itself seems to occur repeatedly between the DBs, across both Africa and South-East Asia, and repeated conversion could mediate maintenance of the shared form (Fig. 5). Given the rest of DBLMSP and DBLMSP2 are very diverged (Fig. 2), either conversion only occurs in the DBL region, or introduction of novel sequence by conversion was only selected in the DBL domain. The latter is perhaps more likely given the biological importance of DBL domains in *Pf*.

In other *Pf* proteins, the DBL domain is key to parasite invasion and persistence. In the EBA family, the DBL domain mediates RBC invasion and in the *vars*, it binds to receptors on epithelial cells and other infected RBCs (iRBC), enabling iRBC sequestration (3,25). Its function in DBLMSP and DBLMSP2 is not known, although one study found it binds to human IgM (27). As an alternative to selection for immune evasion, it could be that the shared DBL domain form mediates binding to a shared environment, in which the two genes are co-expressed, while the private forms mediate binding to private environments. This could be consistent with DBLMSP being expressed in blood-stage, asexual merozoites, and DBLMSP2 being likely only expressed in a subset of schizonts committed to gametocytogenesis (sexual cycle; (44)); gametocytes are reported to preferentially occur in specific niches like the human bone marrow (45). This could perhaps be tested biologically (e.g. using AVEXIS (46,47)), or computationally, using future protein-protein interaction prediction methods based on AlphaFold for example (48).

In conclusion, our study highlights the importance of paralogous gene evolution, and we hope that the higher-resolution sequence data in the DBs will help contextualise their function once it is elucidated.

## Methods

### For results section 1

### Sample preprocessing

Of the >7,000 samples available in the MalariaGEN release that we analysed (released November 2020 (22)), 5,970 were labelled as analysis-ready by the consortium (e.g. after filtering out samples where less than 50% of the genome was callable due to coverage or samples with evidence of more than one infecting species (22)).

The sequenced life stage of *Pf* is haploid, so a single haplotype can be expected in each sample. However, samples with multiple co-occurring strains (MOI > 1) are common in *Pf*, and hard to genotype confidently. We thus further filtered out samples with evidence of MOI> 1, using the *F*_*WS*_ metric (18), which correlates well with MOI in *Pf* (49). We used the *F*_*WS*_ values computed by MalariaGEN (22) and their threshold of > 0.95 for clonality, leaving 3,589 samples for analysis.

### Read preprocessing and quality control

For each sample in the analysis set, reads were downloaded from the ENA, trimmed using trimmomatic (50) to remove adapters and low-quality bases from read ends and subsampled to a maximum of 60-fold expected coverage using rasusa (51). Trimming enables better genotyping with gramtools (see below), and 60-fold coverage is sufficient for genotyping and avoids excess computation. We then characterised the preprocessed reads. Across all 3,589 samples, read lengths had two modes, at 75 and 100 bp, and the estimated per-base sequencing error rate had a single mode of 1×10-4 (supp. fig. M.1, upper panels).

We also aligned reads to the 3D7 reference genome using bwa-mem (52) and measured the average fold-coverage and estimated sequenced fragment lengths (from the distance between aligned paired-end reads). We measured these in alignments across 240kbp of the *Pf* genome defined as ’core genome’ by (31), which excludes highly repeated or variable genomic regions like the telomeres and *var* genes. Fold-coverage mostly lied between 25 and 50 and sequenced fragment lengths between 200 and 300 bp (supp. fig. M.2, lower panels). Some samples had a fold-coverage <10 and were dealt with in a subsequent filtering step described below.

### Pipeline components

In Table 1, we outline the methodology and strengths of each step of our new pipeline. Cortex can assemble large diverged alleles and is highly specific (low false-positive rate), but misses many alleles (low sensitivity). To recover sensitivity, we use Octopus, a pileup-based variant caller (like GATK), but that outperformed GATK in their benchmark, and pays attention to indel-calling in repeat-rich regions (53). We then combine the output of both Octopus and Cortex using our genome-graph-based genotyping tool, gramtools, in a process we call *adjudication* and introduced in (54). The essential idea is to map reads to a graph containing alternate variant calls, and use the mapping of the reads to determine which caller (Cortex or Octopus here) was correct, when they conflict. After adjudication, we run gapfiller, a semi-global assembly-based caller, to recover remaining missing diverged alleles. Gapfiller can efficiently assemble the sequence between aligned read pairs using both reads mapped in the region and all unmapped reads (55).

**Table 1.** Characteristics of the tools used in our new genotyping pipeline. Each tool’s approach and main strengths are summarised. Specific refers to low false-positive rates in variant calling, and Sensitive to high true-positive rates.

| Tool | Method | Strengths | Reference |
| --- | --- | --- | --- |
| Cortex | Global assembly | Specific; can assemble diverged regions | (56) |
| Octopus | Pileup analysis and local assembly | Sensitive and specific; good for SNPs, indels and repeats | (53) |
| gramtools (adjudication) | Genome graph of variation from multiple callers | Leverages different callers' strengths | (54) |
| GapFiller | Sequence assembly between read pairs | Can assemble diverged regions | (55) |
| gramtools (joint genotyping) | Genome graph of population variation | Allows missed call to be found | (28) |

Finally, we perform joint genotyping with gramtools. We built a genome graph containing the variation found in all analysed samples across both genes (plus the 3D7 reference sequence for the rest of the genome), and genotyped each sample using this graph. This enables recovering variants missed in an individual sample, but present in the graph by having been recovered in other samples.

### Genotyping evaluation and performance

We provide only a brief summary of genotyping evaluation and performance here and refer the reader to the supplementary for full details. We used two approaches to evaluate the genotype calls made by both pipelines. The first used 14 independent samples with both Illumina and PacBio data, from which high-quality truth assemblies were built in (31). Using the Illumina data only we genotyped these samples with both pipelines and compared the calls to the truth assemblies. The second relied on remapping the sequencing reads of our 3,589 analysed samples to an ’induced reference genome’, made by applying all the calls made by a pipeline to the 3D7 reference genome. We then measure genotyping quality through the level of agreement between reads and the induced genome. From the read pileups in DBLMSP and DBLMSP2 (appropriately translated from 3D7 coordinates) and for each sample, we measured the number of positions with low coverage, the number of positions where the majority-base in the reads differed from that in the induced reference, and the number of reads with large insert sizes (see supplementary for how we defined large).

We then defined a gene sequence as *confidently-resolved* if, across the full gene (DBLMSP: 2,094 base pairs (bp); DBLMSP2: 2,289 bp), the reads contained no positions with less than 5 aligned reads, no positions where the majority of reads disagreed with the induced reference, and < 15% of reads with large insert sizes. This resulted in 5,895 confidently-resolved DB sequences in total for the gramtools-based pipeline (including the two 3D7-reference representatives). We observed that for the gramtools-based pipeline results, a further 200 DB sequences (92 DBLMSP and 108 DBLMSP2) had no coverage gaps and < 15% reads with large insert sizes, and a single majority-pileup difference. We corrected these single-SNP sequences using a custom script (available with this paper), and added them to our analysis set, and also added the 28 DB sequences from the 14 samples assembled by Otto *et al*. (31), giving a total of 6,123 analysed sequences.

Each step in the gramtools-based pipeline gradually improved genotyping performance across both evaluation metrics, and confidently-resolving most samples at the end (supp. fig. M.3). Overall the gramtools-based pipeline also clearly outperformed the GATK-based pipeline (supp. fig. M.4 and 1.2).

### For results section 2

### Translation to protein

DB gene sequences were translated to protein using seqkit (57), and any resulting sequence with two or more stop codons excluded, as DBLMSP and DBLMSP2 are single-exon genes so that a single stop codon at the end of the sequence is expected (i.e., there is no need to consider stop codons in introns). This removed 3.8% (234/6123; 5,889 analysed sequences left) of the analysis-ready sequences from our gramtools-based pipeline and 8.0% (178/2223) of those from the GATK-based pipeline. Of our 234 removed sequences, 206 samples had one full-length and one truncated protein sequence, while in the remaining 14, both protein sequences were predicted to be truncated. We analysed polymorphism levels at the protein level as, proteins being the functional units of cells, they are more likely to be directly under selection.

### Heterozygosities

At a given aligned position, we define the set of all observed amino acids as {*a*_1_, *a*_*i*_, …, *a*_*n*_ }, and their frequency in the sequences of gene *j* as *f* _*j*_(*a*_*i*_). Then the between-gene heterozygosity for genes *j* and *k* is 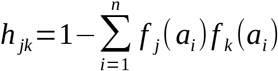

The within-gene heterozygosity is the equation above evaluated on a single gene *j* (*h* _*jj*_). The latter is classically called simply heterozygosity, and is closely related to the ’nucleotide diversity’, π (58).

### DBL domain

The DBL domain was annotated on the 3D7 sequence of DBLMSP by downloading its HMM model from InterPro (https://www.ebi.ac.uk/interpro/entry/pfam/PF05424/) and mapping it using hmmscan from the hmmer suite (59).

### Private and shared peptides

A size-10 peptide was defined as shared if it was seen at least once across both genes (at a given position in the MSA). This definition does not consider geography, meaning, for example, a peptide could be called shared but be seen only once on DBLMSP in a sample from Ghana and once in DBLMSP2 in Cambodia. An alternative is to require a peptide to be found in the same geographical unit (e.g. country) at least once to be considered shared, making them more likely to have a common ancestor and be truly related (as opposed to, for example, convergent mutations). We found both definitions to be essentially equivalent, as most shared peptides by our definition are also found on both genes in at least one country (supp. fig. 2.3).

### HMM logos

To produce HMM logos of the private and shared forms, we first split the original MSA into three separate MSAs, assigning peptides in the original sequences based on private or shared status. For each gene sequence in each sample, each peptide 10-mer was pasted to either the gene’s private MSA, or the shared MSA, adding gaps (’-’ character) where necessary (the sample’s entry in the shared MSA receives gaps for a 10-mer assigned to the private MSA, and vice-versa). We then built one HMM model per MSA using hmmer, and produced logos for visualisation using Skylign (60). In the HMM logos, each letter’s relative height is proportional to its relative frequency in the MSA, and the total height of each stack of letters is proportional to ’information content’, a measure of how different the observed amino acid distribution is from a background expectation (e.g. based on Swiss-prot; though total height is not particularly important here).

### For results section 3

### Identifying recombination

The 5,889 DB protein sequences in the DSR were first clustered at 96% identity using cd-hit (61), producing 35 representative sequences. This allows detection of more distant recombination. To see why, consider a sequence A produced by a recombination of two highly-distinct others, B and C. If another sequence D exists that is one SNP away from A, then A will be aligned full-length to D.

To perform ’mosaic alignment’, MosaicAligner (34) (originally called Tesserae) uses a HMM model parameterised by amino-acid emission probabilities, and transition probabilities (match/indel transitions and donor switches) that must be estimated. Following the original model specification for MosaicAligner (34), the maximum-likelihood (ML) values for all alignment-related parameters were first estimated with the recombination probability ρ set to zero, and ρ was then estimated as the value for which the sum of all target alignments to the panel was maximal. The ML (Viterbi) path for each of the 35 representatives was then obtained given the inferred ML parameter values. From the textual output from MosaicAligner, visual representations of the alignments (e.g. Figure 4 panel b) were produced using custom code available with this paper.

All mosaic alignments included at least one recombination breakpoint. To validate these, we compared the edit distance of each target to its MosaicAligner-inferred donor path with the edit distance to the single closest donor. The former was always smaller than the latter (supplementary fig. 3.1).

### Identifying gene conversion

When comparing the sequences of the DBs in individual samples, we aligned their DNA sequences, as gene conversion occurs at the DNA level, and measured the fraction of identical codons, not nucleotides, to match the protein-level analysis. Notably, codon-level identity is closer than nucleotide-level identity to protein-level identity, though a lower-bound of it as two identical amino acids can be encoded by two different codons.

For identifying samples with evidence of gene conversion, we looked for stretches of identical sequence between the two paralogs on the same genome, for all samples where both DB sequences were ’confidently-resolved’, meaning (as defined above) no coverage gaps or pileup-based differences and no high-levels of large-insert sizes. To rule out the possibility of erroneously attributing sequence sharing to a duplication event in one gene, we further filtered out samples with evidence of a possible gene copy-number variation (CNV; see supp. fig. 3.6). Of the 212 samples with a codon-level identity >0.5, three had a possible CNV, leaving 209 samples all shown in Figure 5.

We further validated 8 samples from each gene conversion cluster in Figure 5, by manually inspecting read coverage levels and insert sizes in IGV (62). We found that coverage levels, insert sizes and read pair orientations were all consistent across DBLMSP, DBLMSP2 and an unrelated gene, AMA1, confirming no duplication of the DBL domain had occurred in these samples.

### For results section 4

The hierarchical clustering tree was built using scipy (63), using the Hamming distance (number of differences) between all unique protein sequences in the DBL-spanning region of the MSA of the DBs (this is equivalent to the edit distance between unaligned sequences). Sequences were clustered by the ’average’ method, i.e. iteratively linking the two clusters with the smallest average distance. This is also called UPGMA. The tree is thus not a ’true’ phylogenetic tree - notably, it does not model the probabilities of different mutations or mutation rate variation across sites.

### For results section 5

For the *Pf* genetic crosses, we used all four publically-available crosses, between strains 3D7 and HB3 (64), HB3 and Dd2 (65), 7G8 and GB4 (66) and 803 and GB4; the raw data are available in (21) and (37) and listed directly in the data availability section. For the ’clone trees’, we used all available data from (7,36) and (36), building tsv files from their supplementary tables and clone tree figures. For six samples, we found convincing evidence of sample mislabeling, confirmed with the original authors. We provide both uncorrected and corrected tables in the repository associated with this paper (see data availability).

We downloaded all available read accessions (284 clone tree samples in six clone trees, and 142 samples in four crosses) from the ENA. For each sample, we performed preprocessing as above (trimmomatic + rasusa) and then genotyped each sample with gramtools on the graph built from the 3,589 ’analysis-set’ samples above. To discover any missed variation (as these samples were not part of the 3,589 in the graph), or mutational events in progeny samples, we then ran all steps of our gramtools-based pipeline up to and including gapfiller (Figure 1).

By our evaluation pipeline standards, all samples were confidently-resolved. We then aligned all pairs of progeny and parent samples (in the clone trees, aligning to the only parent, and in the crosses, aligning to both parents and looking for the closest parent (the parents being usually highly diverged)) to infer any mutations. The only mutational events found were two SNPs, one in each DB gene, in one progeny sample of the cross HB3xDd2 (supp. fig. 5.1).

### Data availability and reproducibility

All code and input data tables are publicly-available on github at https://github.com/iqbal-lab-org/paper_pfalciparum_DBs, and frozen on zenodo at https://doi.org/10.5281/zenodo.7677548. The github repository implements our genotyping pipeline and our analysis of DBLMSP and DBLMSP2 sequences, and contains Snakemake workflows to reproduce all steps, including downloading the input data using input tsv tables. The input tsv files are located under analysis/input_data/sample_lists (including ENA run accessions; see repository README.md file for details), and the main ones are also directly provided on zenodo. All MosaicAligner images are also available on the github repository at plasmo_paralogs/docs/latex_recomb_images/doc.pdf and on zenodo.

All software used and versions are stored in the reproducibility/container folder of the github repository, including a definition file for building a singularity image used by all Snakemake workflows. A copy of this image, called singu.sif, is also available on zenodo.

## Acknowledgements

The authors thank Leah Roberts for reviewing the manuscript, Richard Pearson and Gavin Band for discussions of malaria genomics and Richard Pearson for sharing MalariaGEN data ahead of the pf7 release (67).

## Author Contributions

BL: Conceptualisation, Data Curation, Formal Analysis, Methodology, Investigation, Software, Validation, Visualisation, Writing - original draft, Writing - review & editing SM: Conceptualisation, Investigation ZI: Conceptualisation, Data Curation, Methodology, Supervision, Writing - review & editing

