## Supplementary material for "Gene conversion drives allelic dimorphism in two paralogous surface antigens of the malaria parasite *P. falciparum*" (from another version of this preprint: version 1, 2023-02-27)

### DBLMSP and DBLMSP2

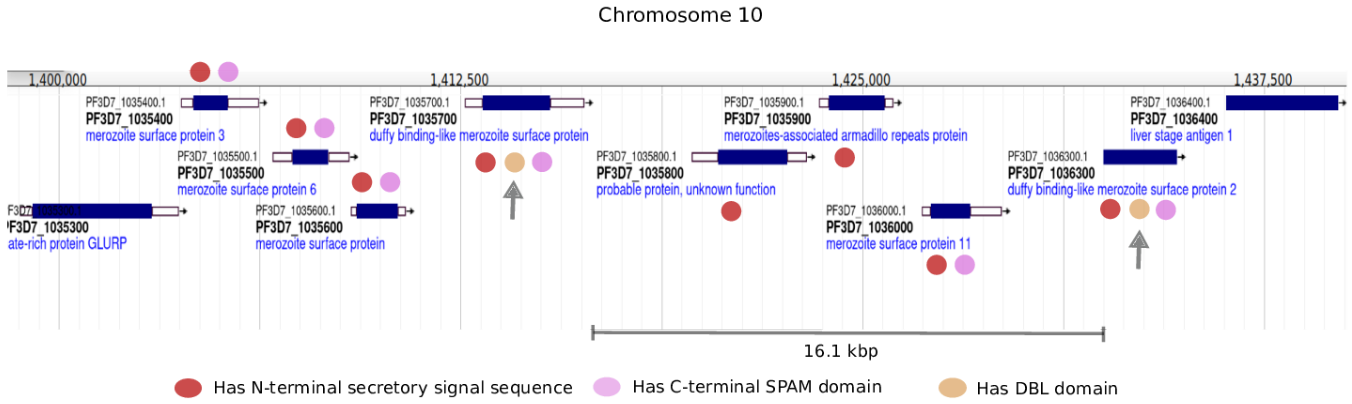


Supplementary Figure 1. **Genomic context and protein domains in DBLMSP and DBLMSP2.** The two genes, marked with grey arrows, are located at a distance of 16.1kbp from each other, inside an array of eight contiguous genes spanning 32kbp on chromosome 10. These genes are likely paralogs due to observed sequence sharing: all eight have an N-terminal shared motif, a further six have a C-terminal SPAM domain, and DBLMSP and DBLMSP2 further share a DBL domain (domains shown as coloured circles below each gene). Figure annotated from a screenshot of the gene track for DBLMSP2 taken from PlasmoDB.

### MalariaGEN data

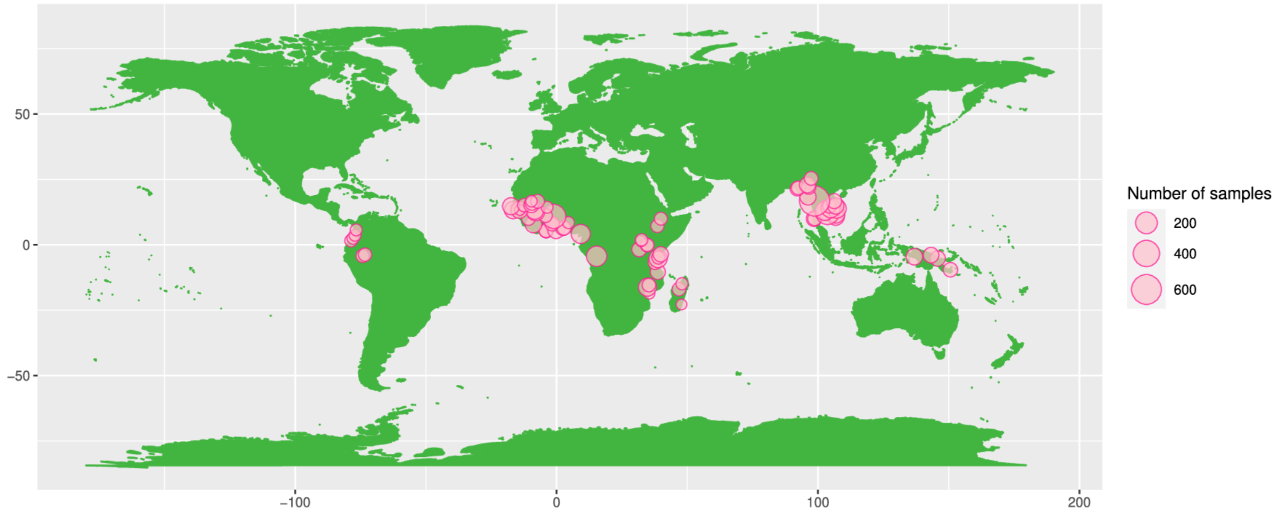


Supplementary Figure 1.1. **Geographical distribution of the 3,589 analysed Pf samples**. A total of 29 countries are represented, with most samples located in the two regions with highest endemicity: Sub-Saharan Africa and South-East Asia.

### Genotyping evaluation framework and performance

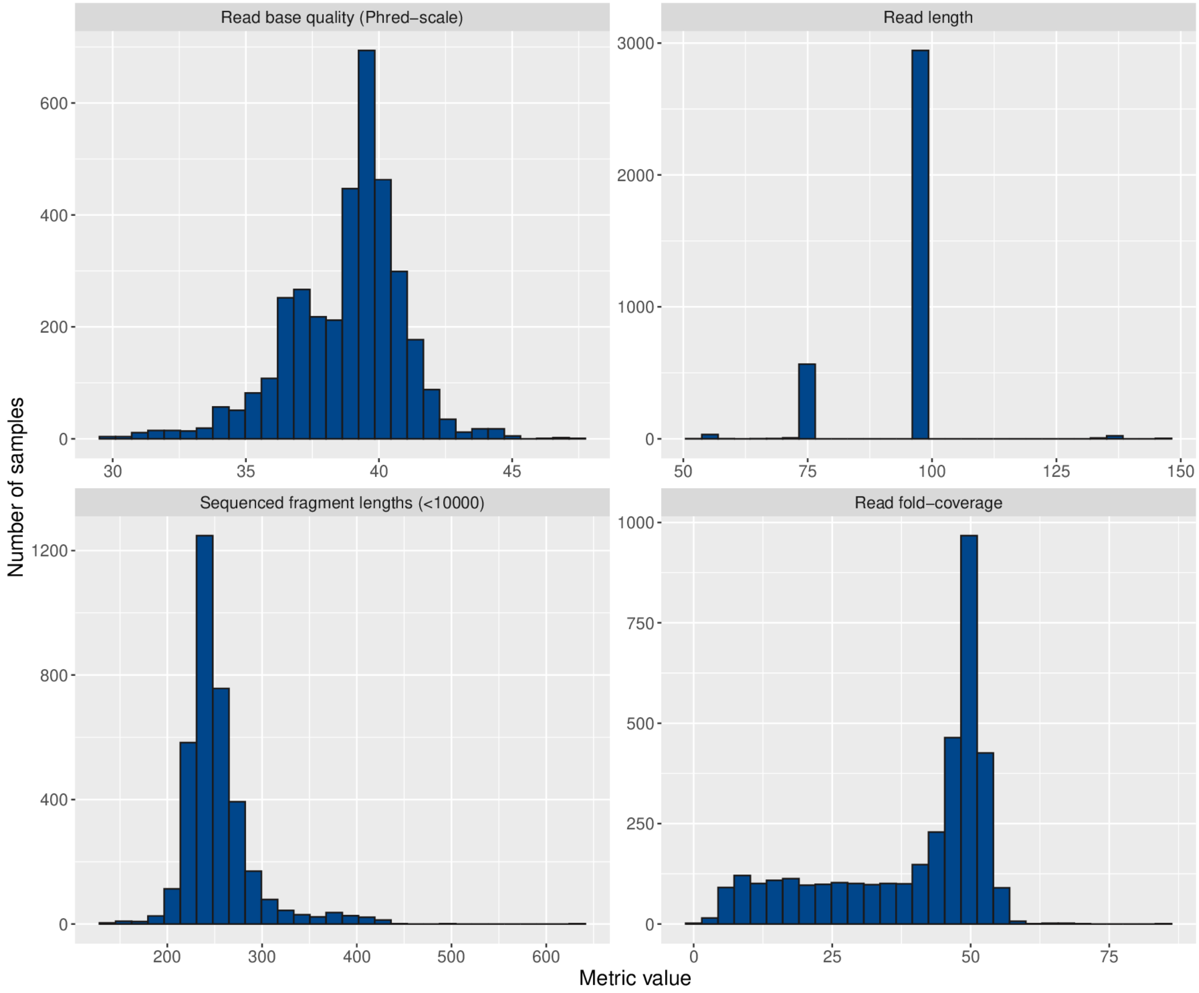


Supplementary. Figure M.1. **Read statistics for the 3,589 analysed Pf samples**. The upper panels show statistics measured on the reads directly: per-base quality (top-left panel) and read lengths (top-right panel). Per-base quality q gives the Illumina-estimated sequencing error rate $\epsilon$ as $\epsilon = 10^{-\frac{q}{10}}$. The lower panels show statistics measured after mapping the reads to the *Pf* 3D7 reference genome: fragment length, estimated from the distance between paired-end reads (bottom-left) and fold-coverage, estimated from the number of reads at each mapped position.

All evaluations started from VCF files of variant calls (69).

We used two different approaches to evaluate variant calls, both illustrated in supp. fig. M.2. The first approach used 14 samples sequenced and assembled by Otto et al. (2018) (31) using both Illumina (MiSeq, 250bp paired-end) and PacBio technologies. We genotyped the samples using the Illumina data only and compared the output to the assemblies, considering them as the true sample genomes. At each step of our pipeline, variant calls made were applied to the 3D7 reference gene sequence and the resulting ‘induced gene sequence’ was mapped to each sample's truth assembly. Performance was measured as the edit distance between the two sequences divided by the gene length (supp. fig. M.2 part a).

A value of zero indicates perfect agreement between the tool's calls and the truth, while values >0 indicate missing and/or false positive calls. Note that the 14 samples were not part of the 3,589 analysed samples, and thus did not figure in the genome graph used for the joint genotyping step.

To evaluate genotyping calls in the 3,589 analysed samples themselves - i.e., in the absence of a source of independent truth - we used a read realignment-based approach. At each pipeline step, called variants were applied to the *Pf* reference genome to create an ‘induced reference genome’  and the reads remapped to it. Missing or incorrect variants can be identified by looking for inconsistencies between the aligned reads pileups and the induced reference (supp. fig. M.2 part b). Specifically, we measured for each gene the number of positions where the majority of reads disagree with the reference base, the number of positions with no (or low) read coverage, and the number of read pairs with abnormal insert sizes (i.e. falling outside the bulk of the size distribution).

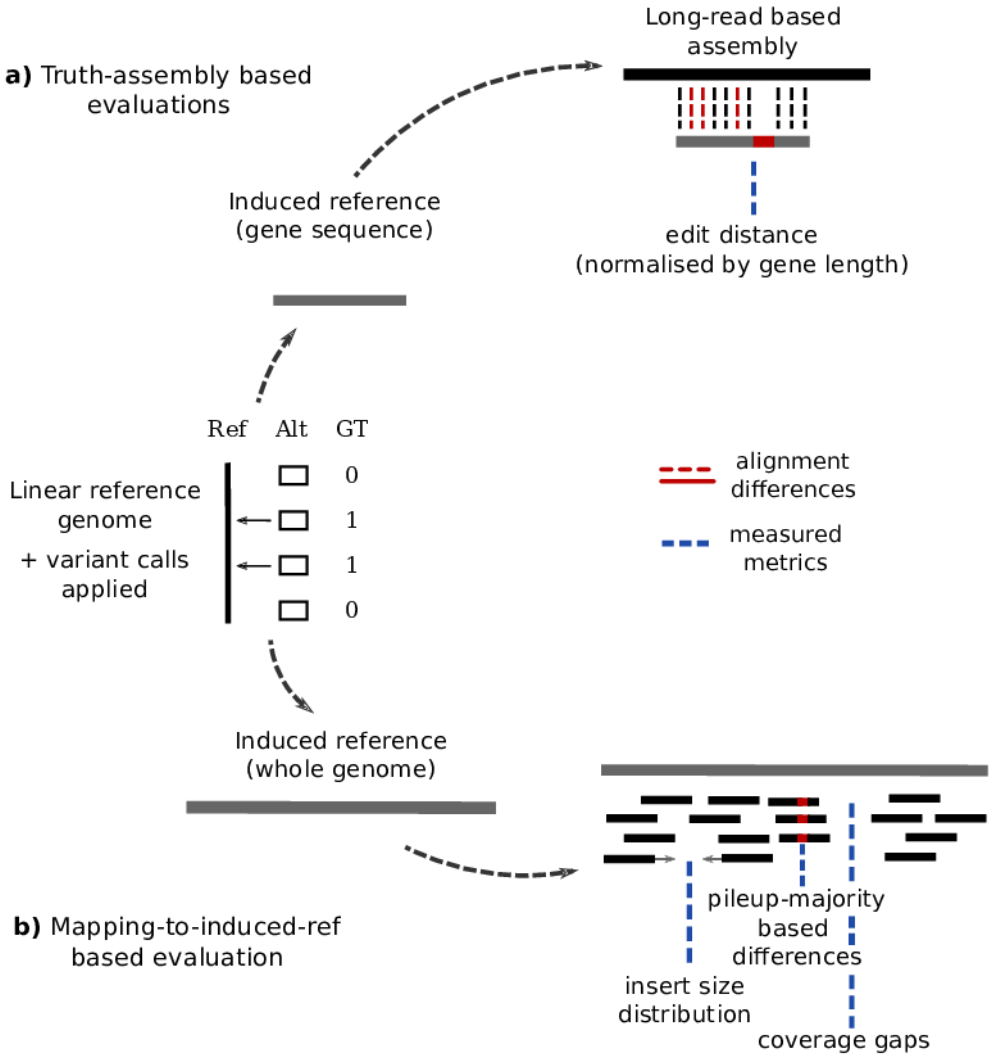


Supplementary Figure M.2. **Framework used to evaluate the variant calls from the GATK and gramtools-based pipelines**. Starting from a tool's variant calls in a VCF file (middle), two independent evaluations were performed. First, for 14 samples with truth assemblies, the calls were directly compared to the truth, by applying them to the 3D7 gene sequence and measuring edit distance for the whole gene (part a). Second, the calls were all applied to the reference genome, and the reads remapped to this induced reference. Incorrect or missing calls then appear from read pileups, as majority-differences compared to the reference base, coverage gaps, or inconsistent insert sizes between read pairs (part b).

In supp. fig. M.3, we show the evaluation results for each step of the gramtools-based pipeline in both DBLMSP and DBLSMP2. Panels a and b correspond to those in supp. fig. M.2. Across both panels, and for both genes, each step of the genotyping pipeline improves (or is at least as good as) the previous one, and all metrics approach zero after the final step ('gram_joint_geno', beige colour). In panel b, only a subset of 500 samples was evaluated, selected to be as geographically wide-ranging as possible, to save on compute and storage requirements (406GB disk size for 500 compared to 2.8TB for all samples). For comparing our pipeline and the MalariaGEN results (see below), all samples were evaluated.

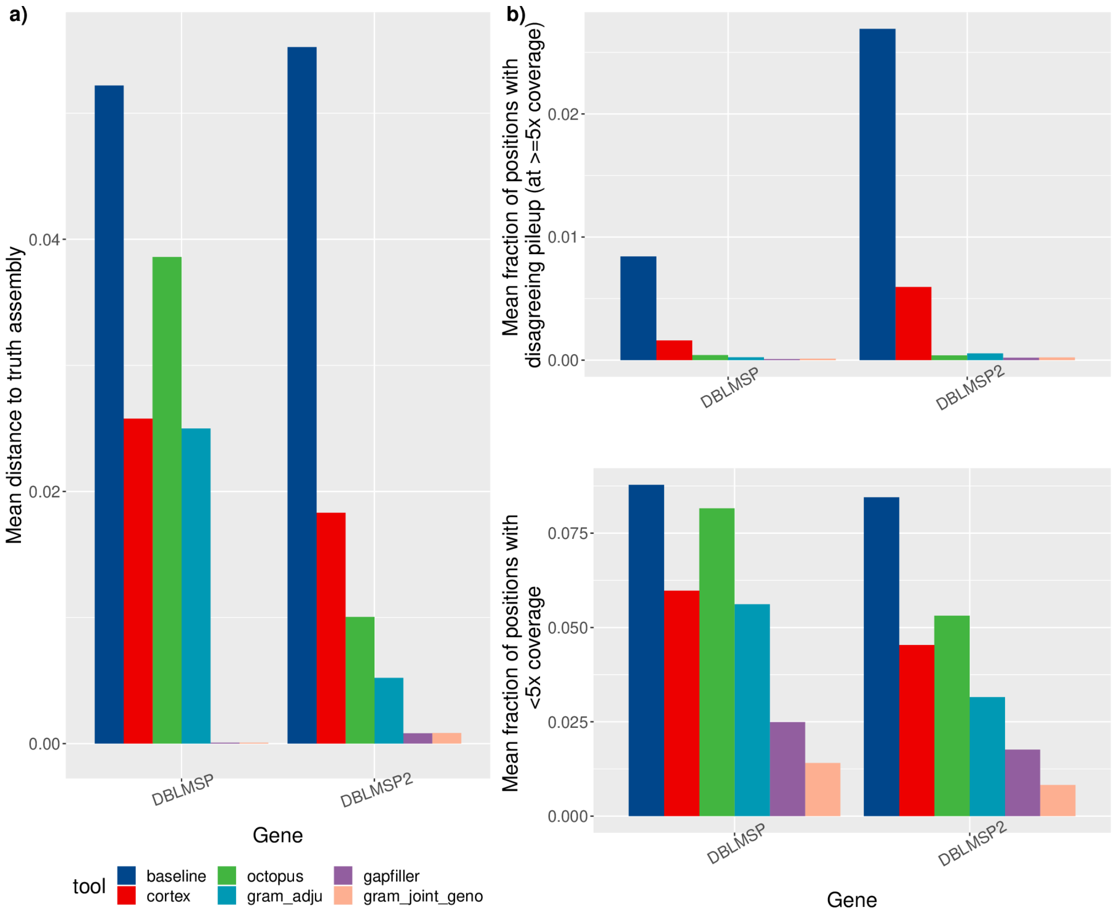


Supplementary Figure M.3. **Performance of the gramtools pipeline steps in DBLMSP and DBLMSP2**. The two panels, a and b, correspond to parts a and b of the evaluation framework in Fig. 7. Panel a shows the mean edit distance between the inferred gene sequence and the truth assembly for the 14 samples with truth assemblies (edit distance is scaled by gene length). Panel b shows the fraction of positions with pileup-based differences (top) and with low read coverage (bottom), after the sequencing reads are remapped to the 3D7 reference genome with each tool's called variants applied. A pileup-based difference is when the majority of reads disagree with the reference at a given position, given a minimum of five mapped reads, and low read coverage is defined as a position with fewer than five mapped reads. Each bar in panel b shows the mean across 500 of the 3,589 analysed samples. Across both panels, each coloured bar corresponds to one additional step in the gramtools-based pipeline, in the same order they are run (see Fig. 1). The ‘baseline' condition is not part of the pipeline and refers to using 3D7 reference gene sequence with no variants applied (in panel a: 3D7 sequence aligned to the truth assemblies; in panel b: sample reads aligned to the 3D7 reference genome).

In supp. fig. M.4, we compare the results of the final step of the gramtools-based pipeline with those from the GATK-based pipeline, showing the same metrics as in supp. fig. M.3. For GATK, results on the 3,589 analysed samples (panel b) were taken from the 2021 release (22), while for the 14 truth assemblies (panel a) they were taken from the 2023 release (68). For both genes, our pipeline clearly outperforms the GATK-based approach on all metrics.

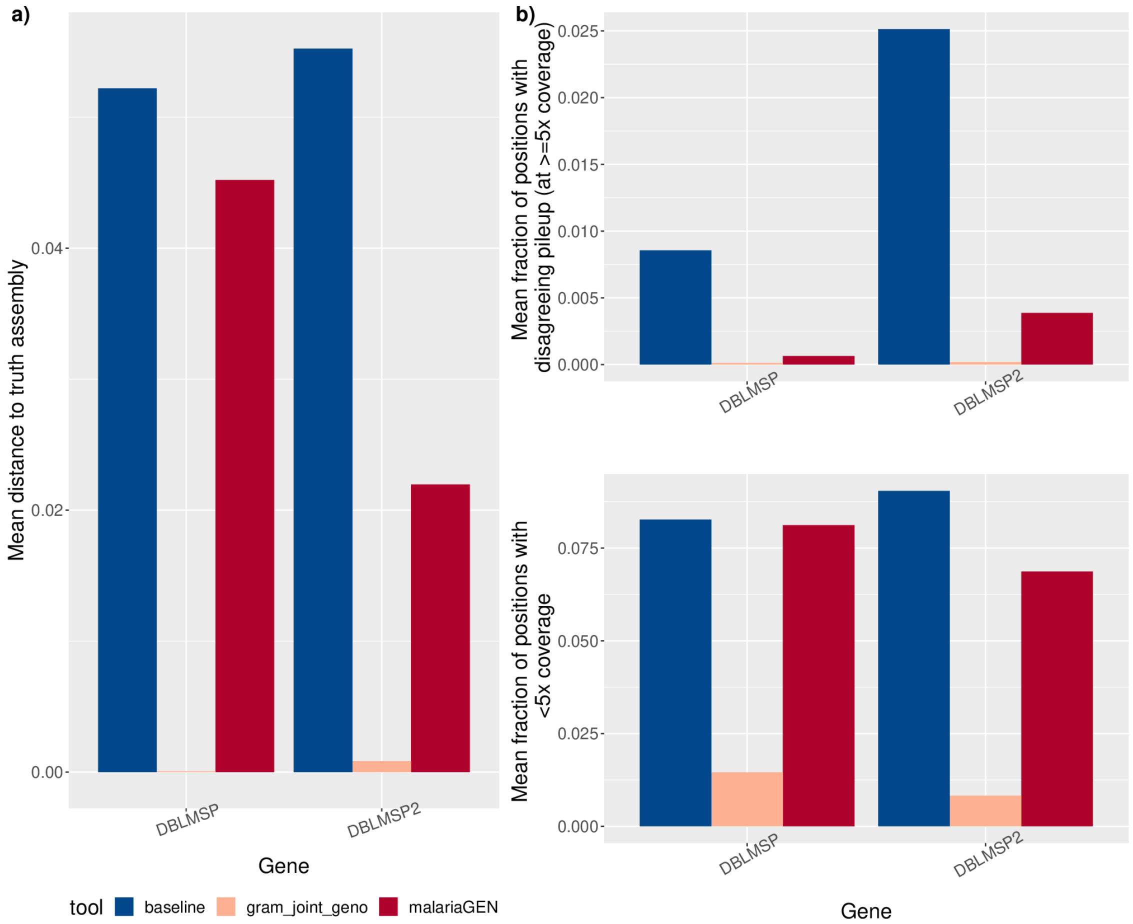


Supplementary Figure M.4. **Global performance of the gramtools-based and GATK-based pipelines**. Panels a and b show the same metrics as supp. fig. M.3. Metrics in panel b were computed on all 3,589 analysed samples.

In supp. fig. M.3 and M.4, only the mean values across samples are shown. We show the full frequency distribution for the data in supp. fig. M.4 panel b in supp. figure M.5.

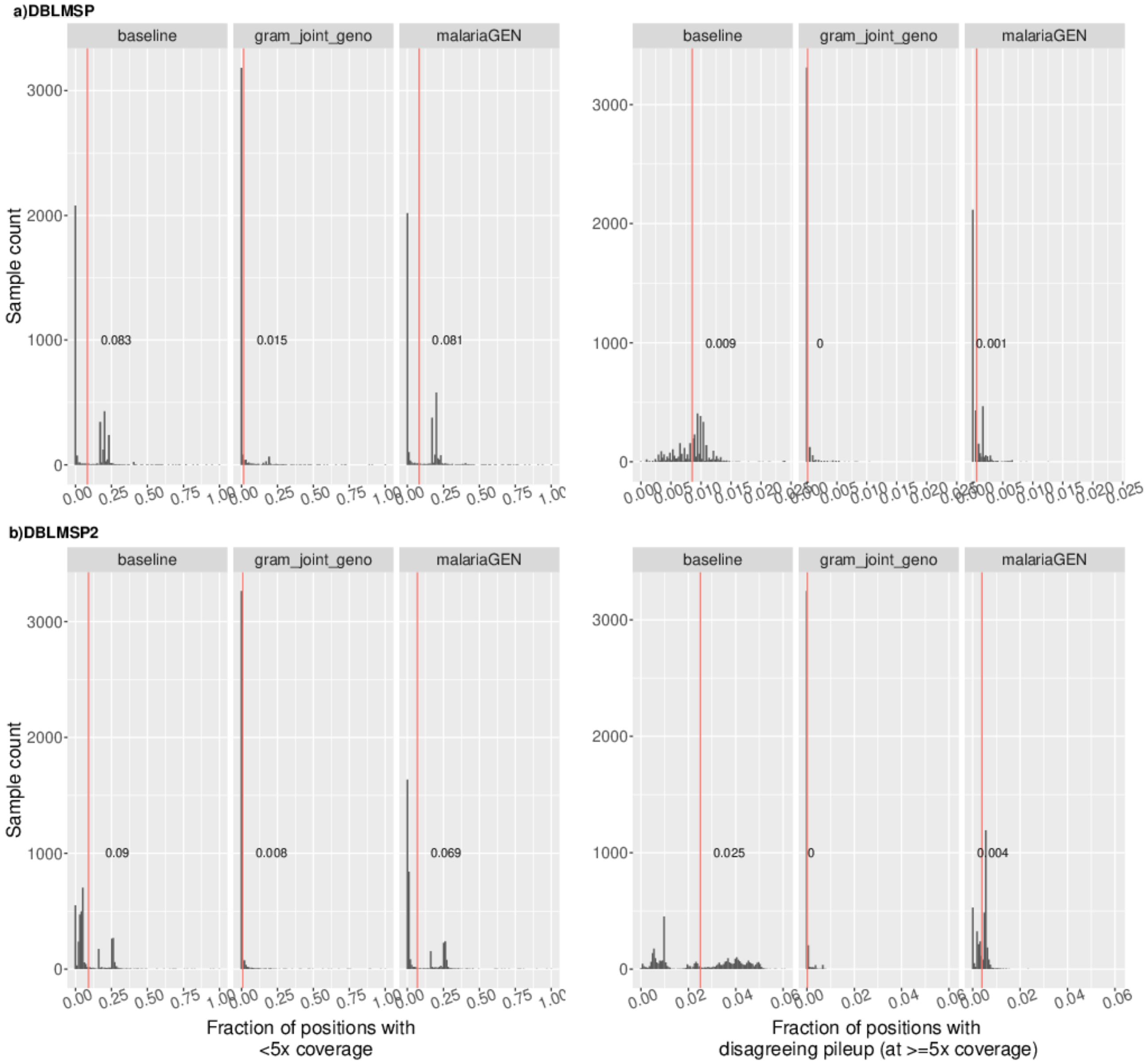


Supplementary Figure M.5. **Frequency distributions of the evaluation metrics for the gramtools-based and GATK-based pipelines.** Each subplot shows the frequency distribution, across all 3,589 analysed samples, of the fraction of positions with pileup-based gaps (left-hand side plots) or differences (right-hand side plots), for DBLMSP (top) and DBLMSP2 (bottom). The mean is shown as a red vertical line (value shown in text next to it) and corresponds to the height of the coloured bars in Fig. M.4 panel b.

Based on the realignment metrics computed on the 3,589 analysed samples (supp. fig. M.4 panel b), we designed a set of criteria to produce a final set of DB sequences for analysis. To consider a sequence for downstream analysis, we required three conditions to hold: i) none of the gene positions contained fewer than five aligned reads (pileup-based gaps); ii) none of the gene positions contained pileup-based differences, defined as a position where the majority of aligned bases differed from the induced reference; iii) the fraction of reads with large insert sizes was low, where an insert size was defined as large if it exceeded the mean insert size + two standard deviations. We defined ‘large fraction’ as 15% by looking at the distribution of the fraction of large insert sizes across the 3,589 analysis samples in a gene that is confidently-resolved by both pipelines (AMA1), and taking the value below which almost all samples lie.

The first two filters ensured that the remapped reads fully covered and agreed with the inferred gene sequence, and also removed samples with low-coverage, due for example to DNA amplification bias. The last filter ensured that large structural variations were not missed: for example, translocations or duplications can lead to abnormally large insert sizes. Supplementary figure 1.2 shows the number of sequences left after each filter is successively applied, across both DBLMSP and DBLMSP2 (x-axis). The three vertical panels show a ‘baseline’ approach of making no calls, and the two genotyping pipelines (‘gram_joint_geno’, ‘malariaGEN’).

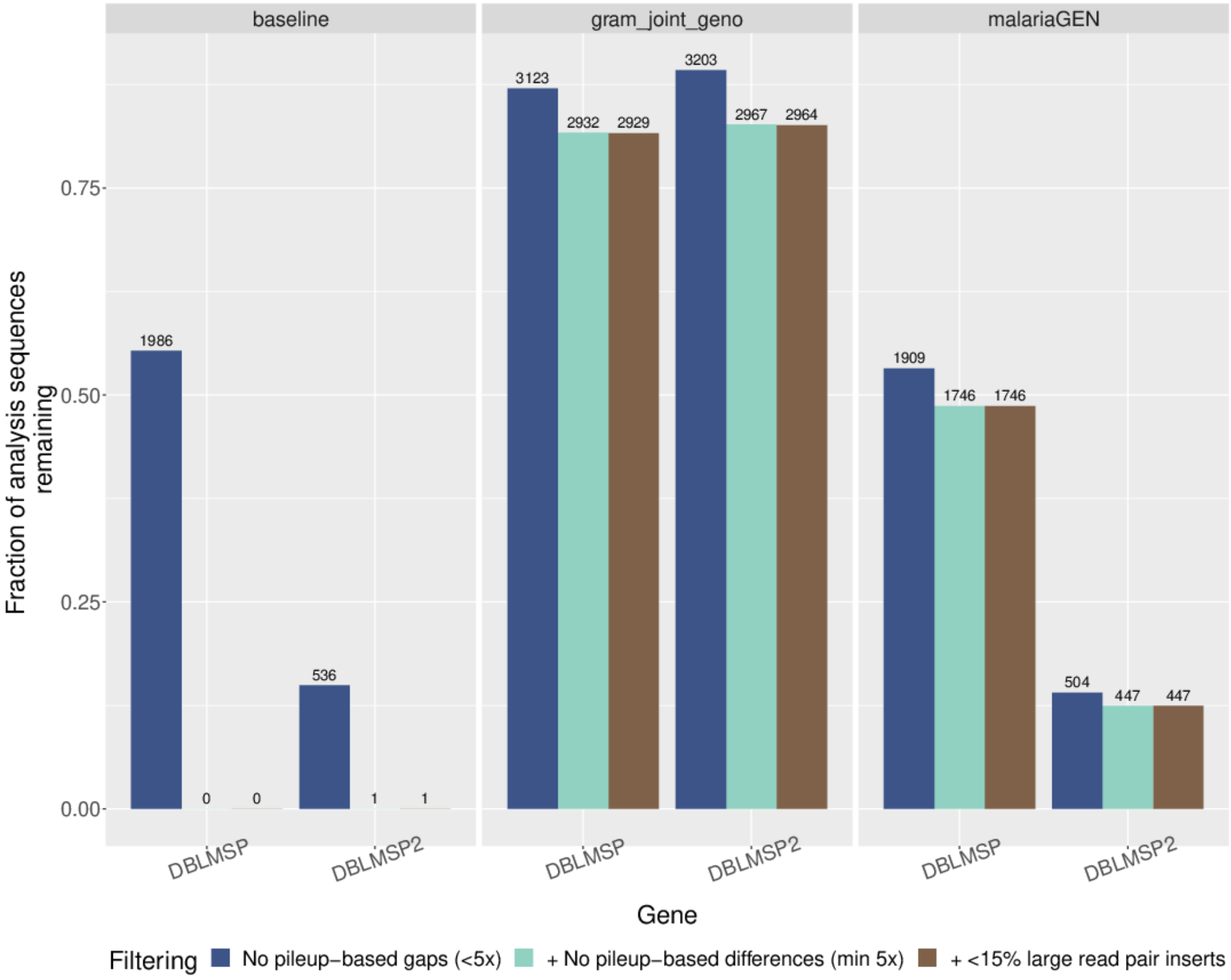


Supplementary Figure 1.2. **Sequence filtering using pileup-based metrics.** In each panel (‘baseline’:no variant calling, ‘gram_joint_geno’: gramtools-based pipeline, ‘malariaGEN’: GATK-based pipeline), the total fraction of remaining gene sequences (out of the 3,589 analysed samples) passing filters is shown. Filters (colours) are applied in succession, on each set of remaining gene sequences, in the order they appear in the legend. The number of remaining sequences is given above each coloured bar.

In total, 5,895 DB sequences from our pipeline met all three criteria (5,893 from supp. fig. 1.2 above (2,929 DBLMSP and 2,964 DBLMSP2 sequences), plus the two 3D7 reference sequences). For an additional 200 DB sequences, criteria i) and iii) were met, but a single pileup-based difference remained in the remapped reads, with a clear single base differing from the induced reference at a frequency >0.5 in the reads. We corrected these single SNPs with a custom script (available with this paper, see data availability) and added these sequences to our analysis set. We also added the 28 DB sequences from the 14 samples with truth assemblies by Otto et al. (31), giving a total of 6,123 analysed sequences.

### Sequence sharing patterns

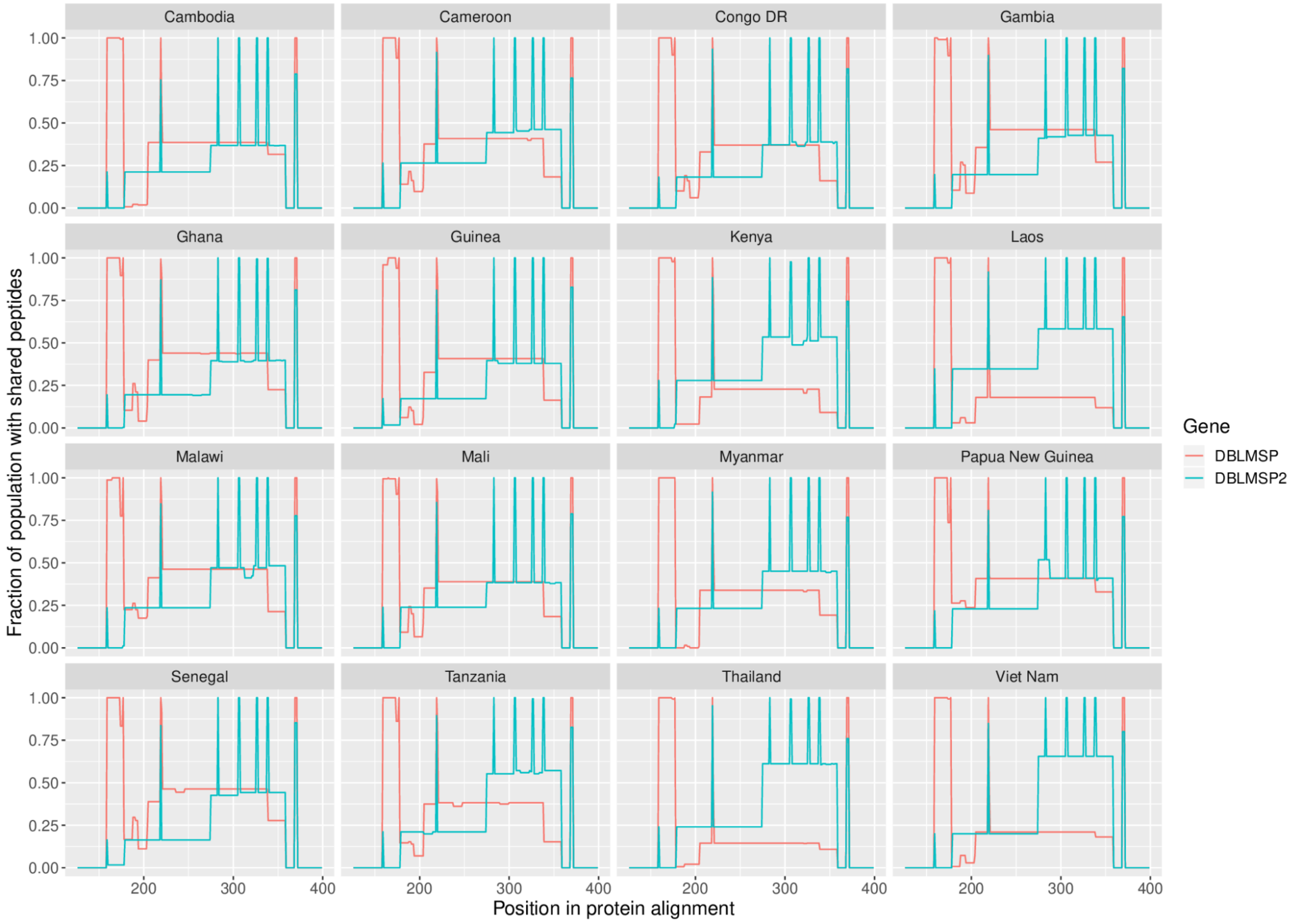


Supplementary Figure 2.1. **Shared peptide 10-mers are highly prevalent.** The frequency of shared peptides (y-axis), at each position (x-axis), is shown for the 16 countries with more than 50 sequences. Colour indicates frequency in each gene. Shared peptides are found at high frequencies, between 25 and 50%, across all countries. By extension, private peptides are also frequent, as any non-shared peptide is a private peptide. Values of zero, at the left and right-ends of the x-axis, show the diverged flanks of the region, while values of one correspond to peptides that are always identical in both genes, i.e. where any mutations are likely eliminated by selection. On the left-hand side of the plots, DBLMSP2 displays a region with low shared peptide frequency across all countries, indicating this region has almost fully diverged between the DBs.

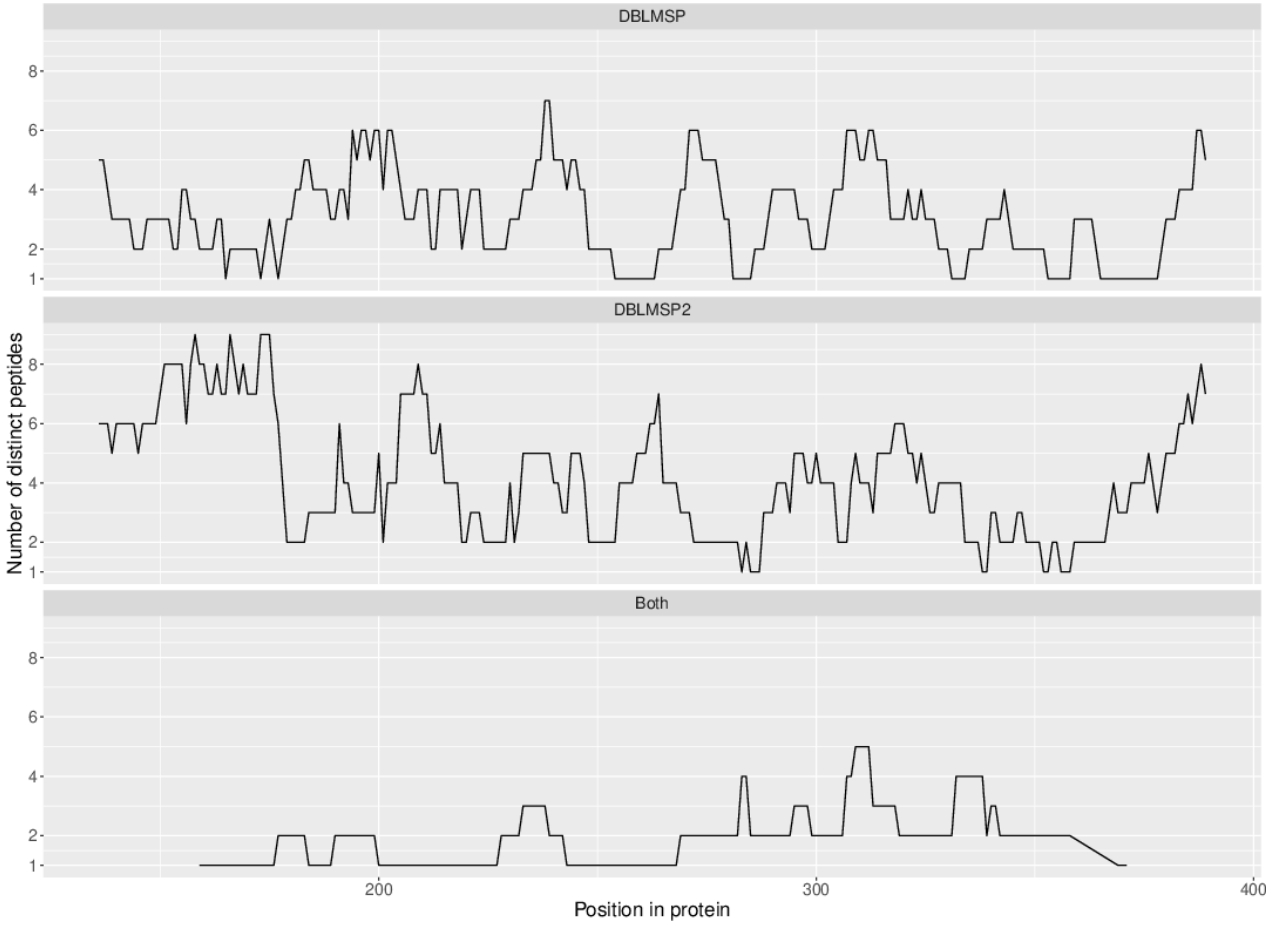


Supplementary Figure 2.2. **Number of distinct shared and private peptides per position**. For the two private and one shared MSAs, containing peptides only found in DBLMSP (top panel), only found in DBLMSP2 (middle panel), or both (lower panel), the total number of distinct peptide 10-mers at each position is shown. Mostly one to four peptides were observed at each position in the shared category, while two to six were observed on each gene only. This figure complements Figure 3, which shows that mostly two 10-mer peptides occur in each gene with high frequency - here, total number is shown, regardless of frequency.

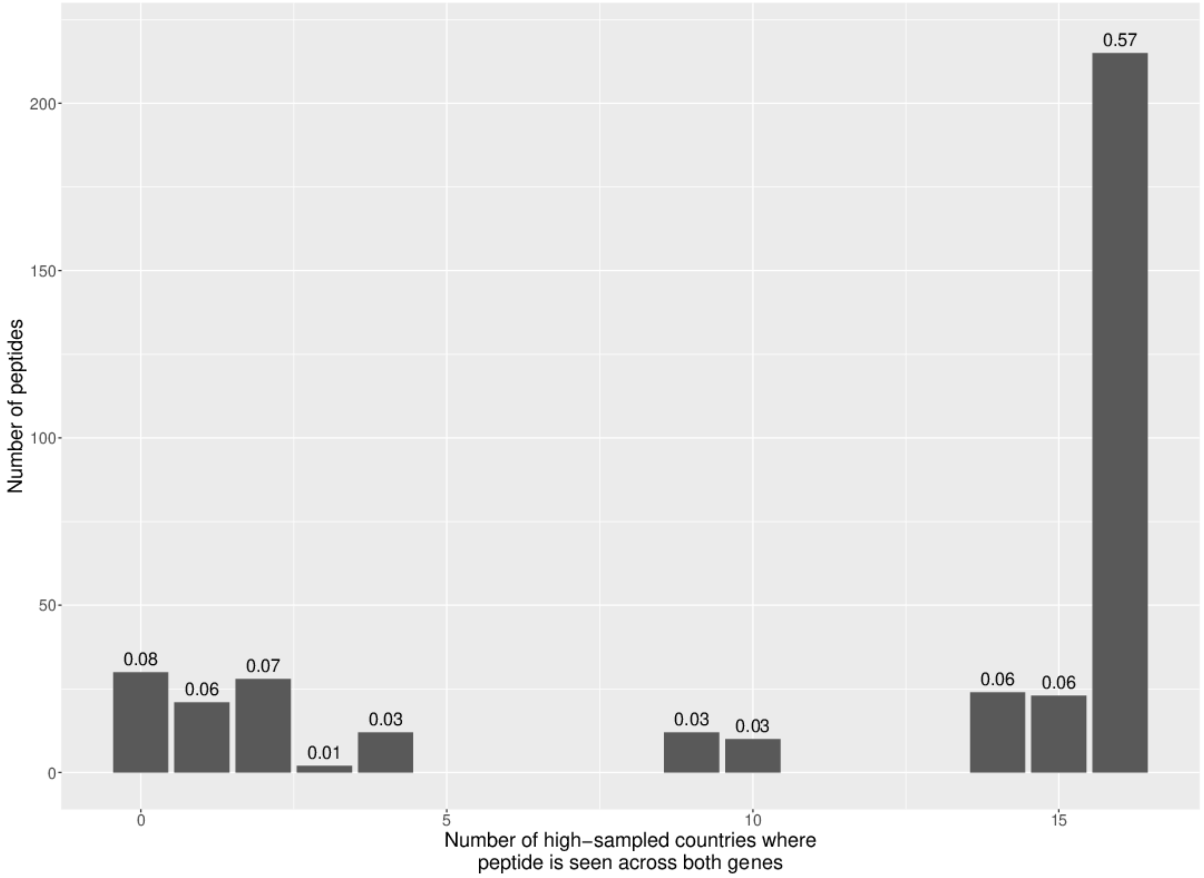


Supplementary Figure 2.3. **Peptides defined as shared are shared in many different countries**. The number of shared peptides by our definition (y-axis) that are found in both genes inside the same country, for up to 16 countries with high levels of sampling (defined as >50 available DB sequences; x-axis). A value of zero on the x-axis means the shared peptide is not found on both genes in any of these countries, and 16 means it is found on both genes in all of them. A majority (57%) of shared peptides are found in all of these countries, and 86% are found in at least two different countries, showing that the shared peptides are, overall, highly widespread geographically.

### Recombination and gene conversion

#### *Mosaic alignments*

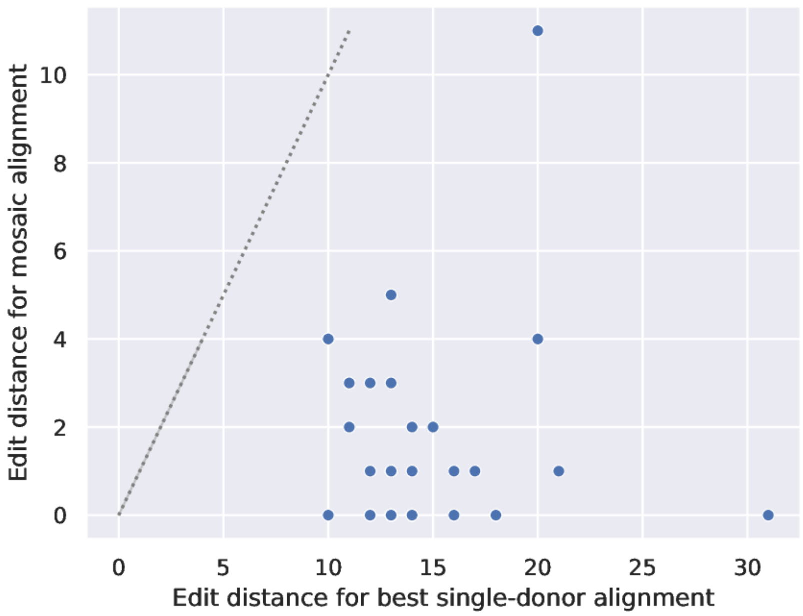


Supplementary Figure 3.1. **Validation of MosaicAligner-inferred recombination events**. The blue dots show, for each target, its edit distance to the path of donors inferred by MosaicAligner (y-axis) and the edit distance to the single closest donor (x-axis). The grey dotted line shows y=x. All inferences reduce edit distances to the single closest donor, supporting adding recombination breakpoints to the alignment.

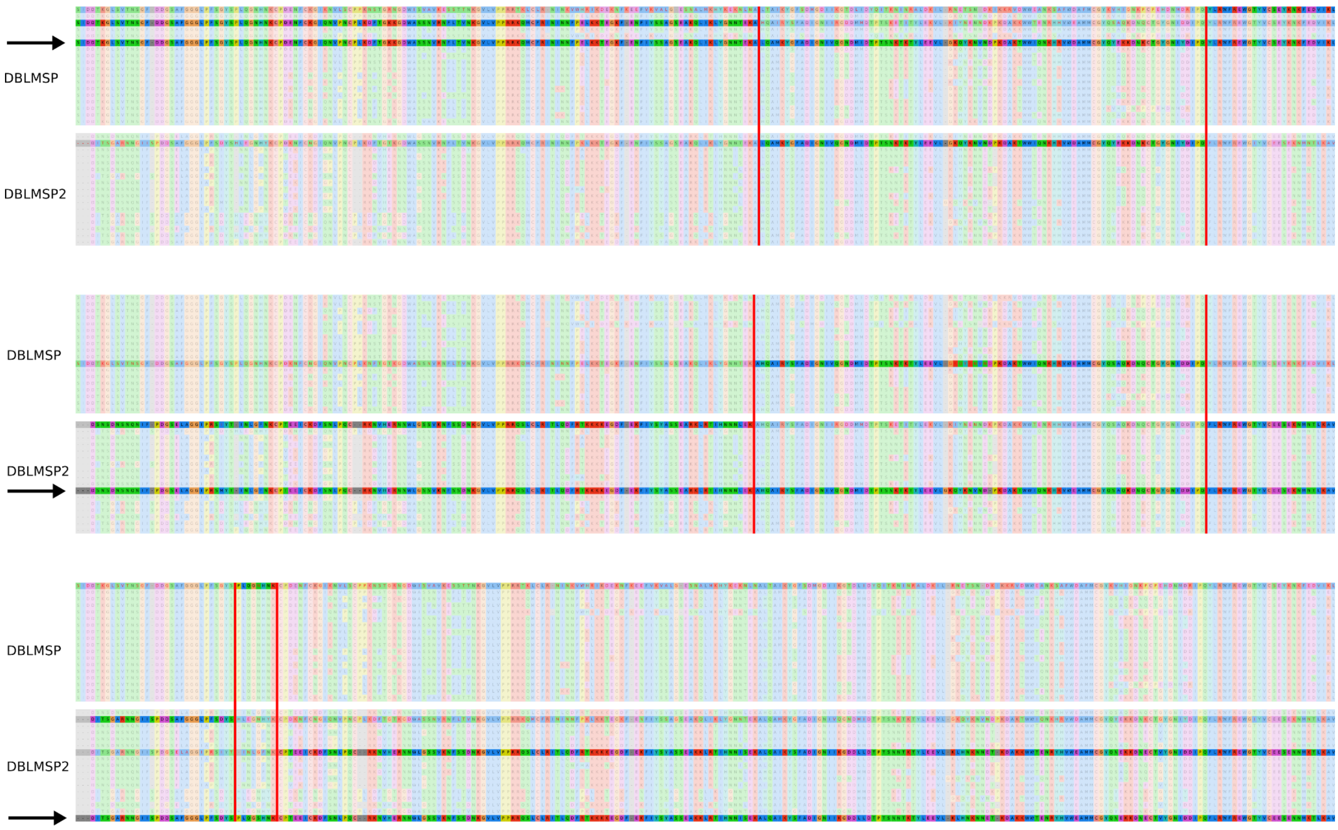


Supplementary Figure 3.2. **Three mosaic alignments support gene conversion.** In each panel, two recombination breakpoints can be seen (red vertical lines). The target sequence is the fully-opaque one (along its entire length; indicated with a black arrow), and the donor sequences (those the target aligns to) are shown as highlighted in places they match the target, and less opaque where they do not. In each panel, the target aligns to donors across the two different genes, consistent with gene conversion between the genes.

*Gene conversion*

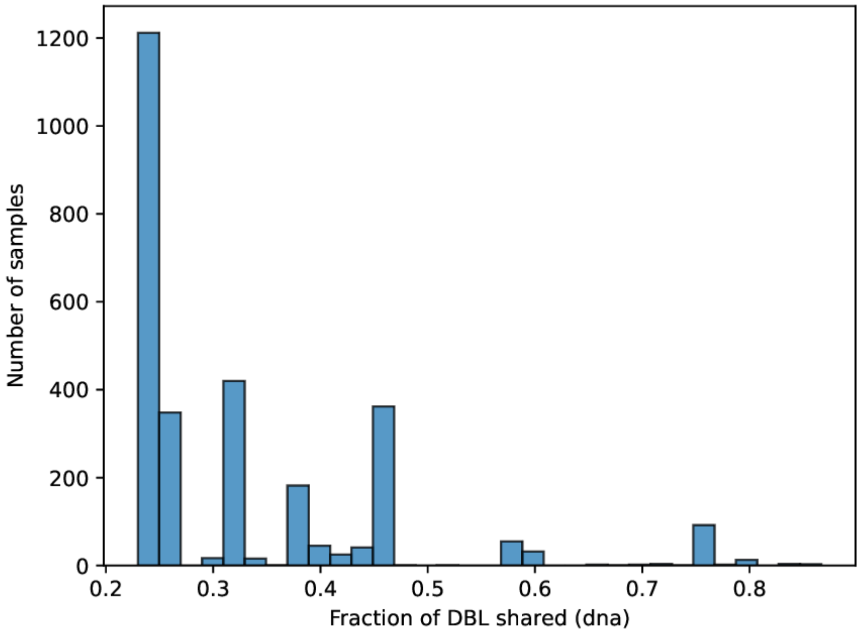


Supplementary Figure 3.3. **Distribution of DB codon-level identity across 2,882 confidently-resolved samples.** For all samples in which both DB sequences were confidently-resolved, the DNA sequences of DBLMSP and DBLMSP2 within a single genome were aligned and the fraction of identical codons in the DSR recorded. Most samples have quite low identity levels (e.g. 0.2 up to 0.4), and a minority of samples have high identity levels, defined as >0.5 identity. The latter samples are illustrated in Figure 5.

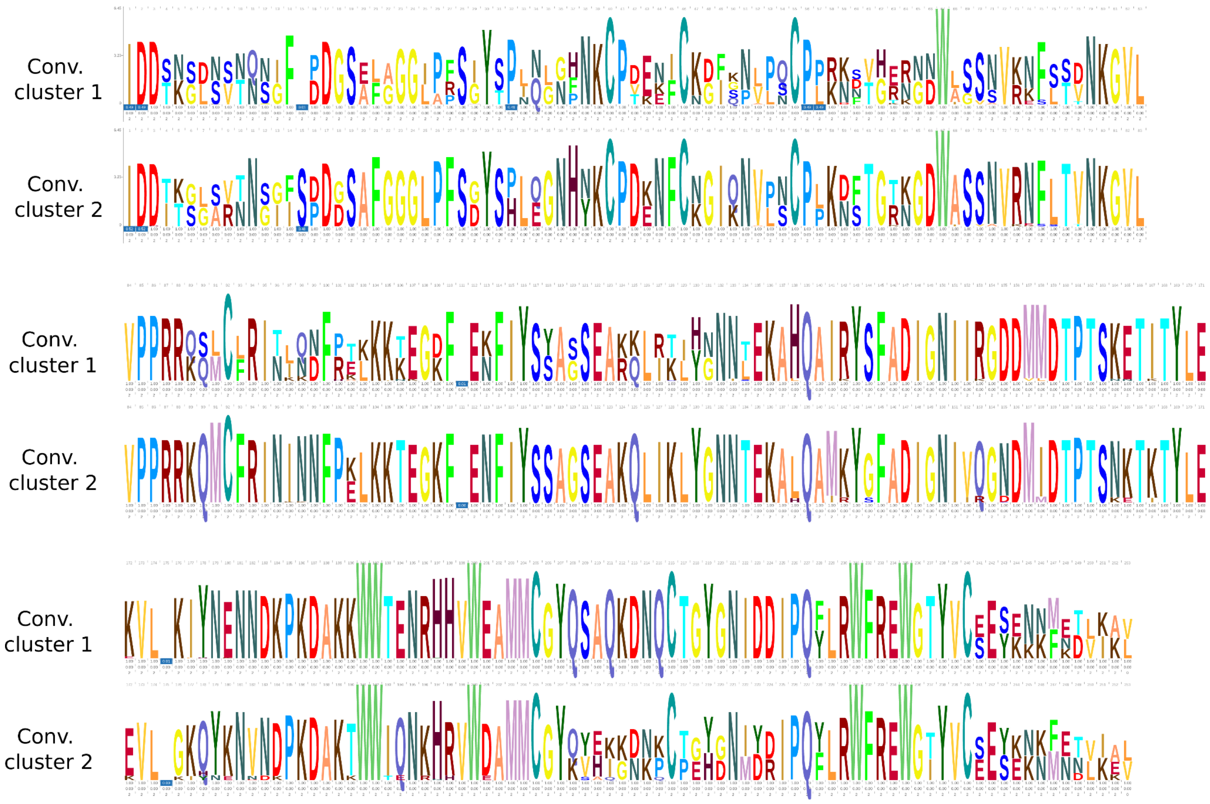


Supplementary Figure 3.4. **Sequence motifs of the two gene conversion clusters.** One logo was produced for each of the two conversion clusters in Figure 5, and each logo split into three portions for visual clarity. While at many positions, the sequences in each conversion cluster overlap, each cluster is enriched for different amino acids, and some positions have entirely different amino acids. This supports a distinct evolutionary trajectory for each cluster and thus at least two distinct gene conversion events having occurred in the DBs.

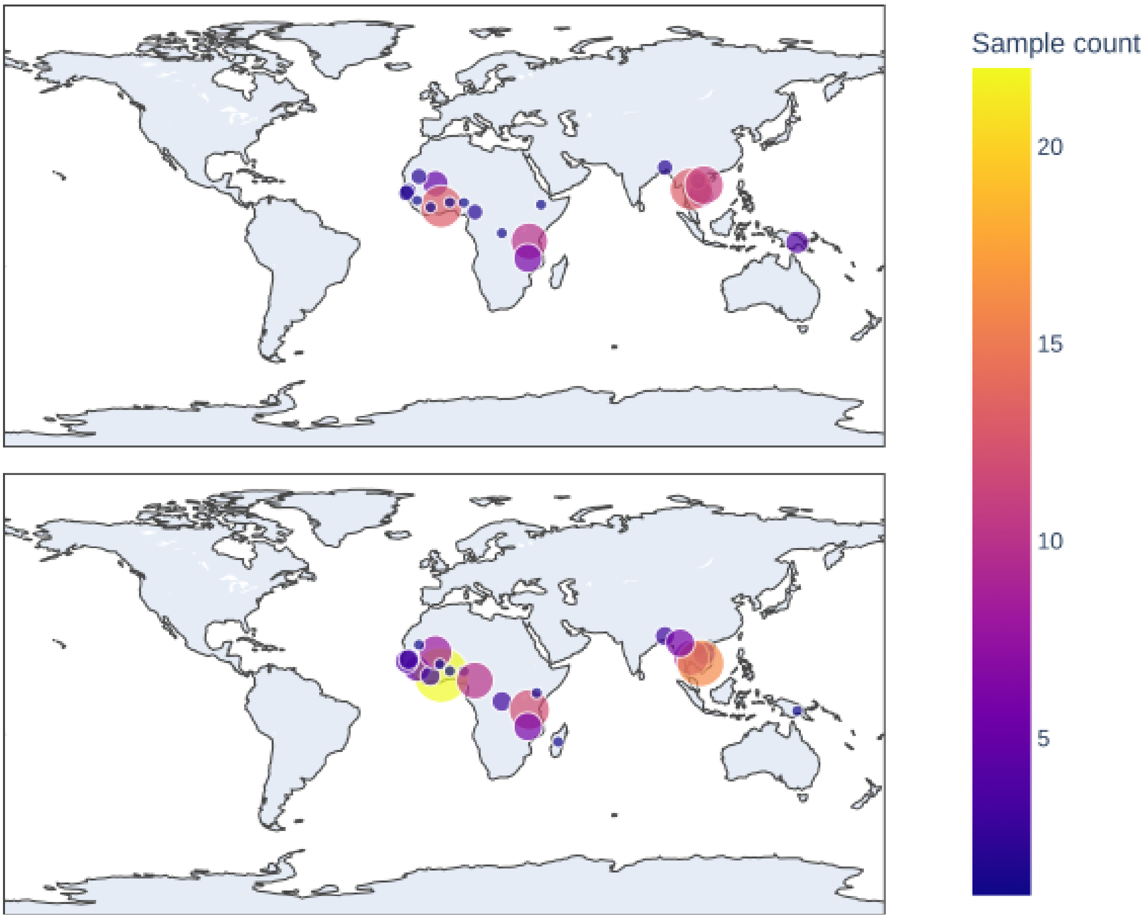


Supplementary Figure 3.5. **Geographical distribution of the two gene conversion clusters.** The two panels correspond to the two gene conversion clusters identified in Figure 5 (in the same order, top to bottom). In each panel, the number of samples in each geographical region is shown, both through the size and colour of dots. For both conversion clusters, samples are geographically widespread, occurring across west and east Africa and South-East Asia.

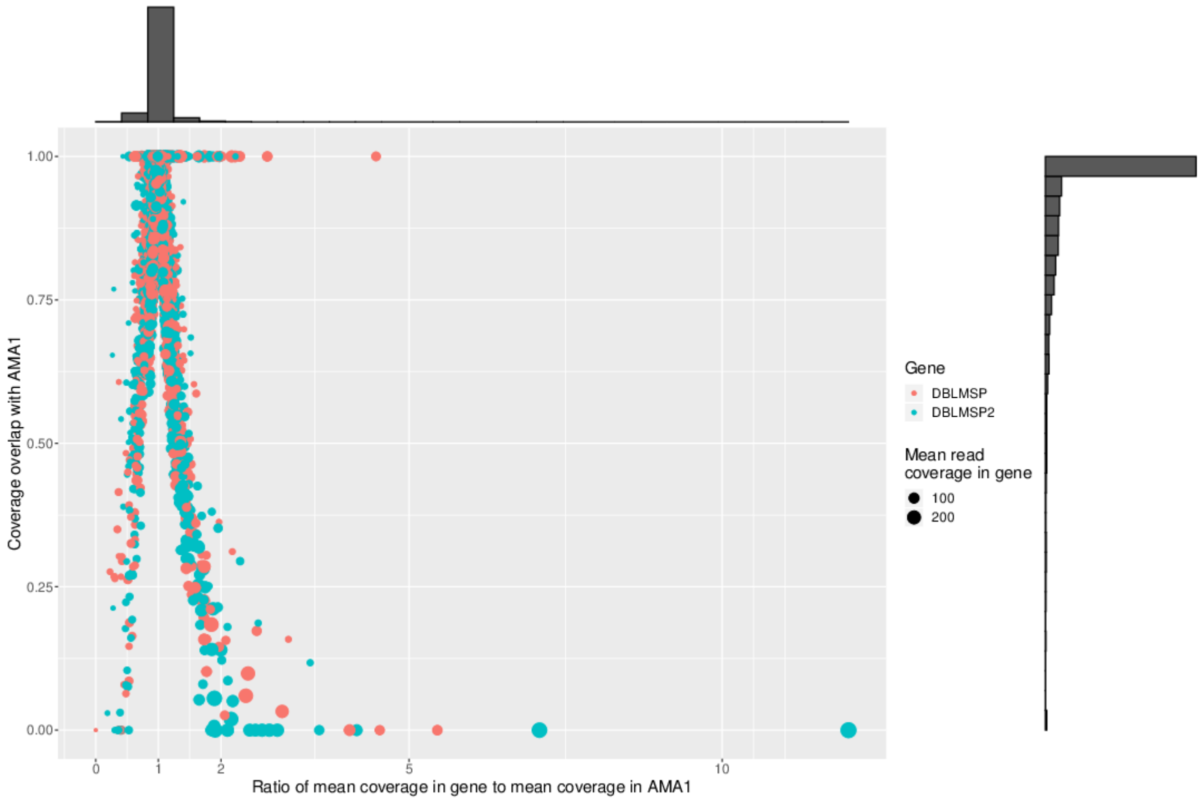


Supplementary Figure 3.6. **Identification of samples with putative CNVs.** For all 3,589 analysis-set samples, the mean and standard deviation (std) of the per-base read coverage of reads realigned the ‘induced reference’ (supp. fig. x, panel b) was measured in genes DBLMSP, DBLMSP2 and AMA1. For each gene, we produced a *coverage interval* {mean - 2*std, mean + 2*std}, that we consider a ‘plausible range’ of gene-level coverage. The x-axis shows the ratio of the mean coverage in the DBs to that in AMA1, a gene that we assume to be single-copy in all samples. The marginal distribution histogram is shown on top. Most samples have a ratio of one, and some have ratios < 0.5 or > 2, indicating possible copy-number changes. The y-axis shows the fraction of the DBLMSP or DBLMSP2 coverage interval overlapped by the AMA1 coverage interval. Most samples have totally-overlapping intervals (marginal distribution on right-hand side). Small overlap values indicate more-likely true differences in coverage. Of the 6,123 analysed DBLMSP and DBLMSP2 sequences (see supp. fig. x), 31 had a fold-coverage > 2 and an overlap < 0.5 (bottom-right of plot), indicating putative duplication. Three of these overlapped with samples with evidence of gene conversion, and were filtered out in that analysis (Figure 5).

Recombination

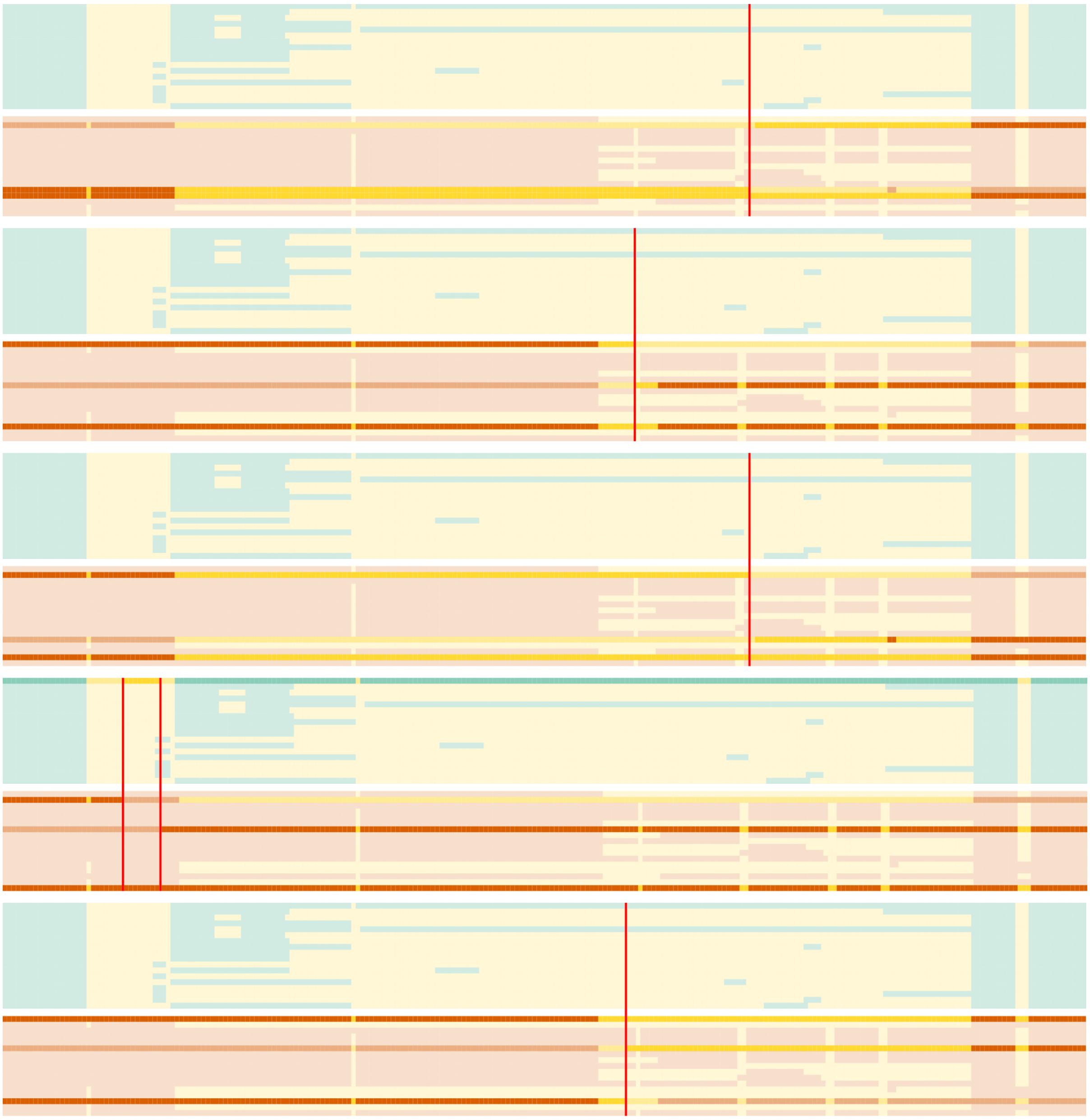

Supplementary Figure 4.1. **Within and inter-form recombinations**. Five different recombinations are shown inside matrices, where, as in fig. 3.1, each matrix depicts the mosaic alignment of one target sequence to the panel of 35 sequences. Sequences from DBLMSP (top) and DBLMSP2 (bottom) are separated by a white horizontal strip. Each cell is coloured by whether the size-10 peptide centred at that position occurs only in DBLMSP (blue-green), only in DBLMSP2 (orange), or in both (yellow). Recombinations mostly occur within the private DBLMSP2 form (all donors are mostly orange) and within the shared DBLMSP2 form (all donors are mostly yellow). In the last panel, the target is a recombinant of a highly-private and a highly-shared sequence.

### Direct evolution in the DBs

**a)** SNP in DBLMSP2

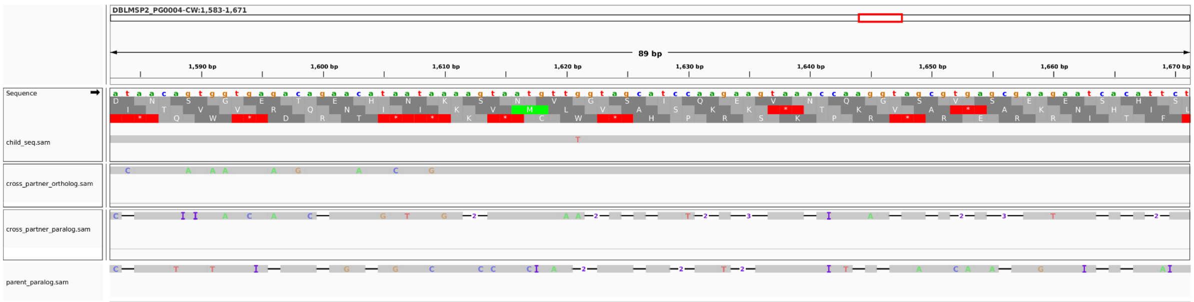


**b)** SNP in DBLMSP

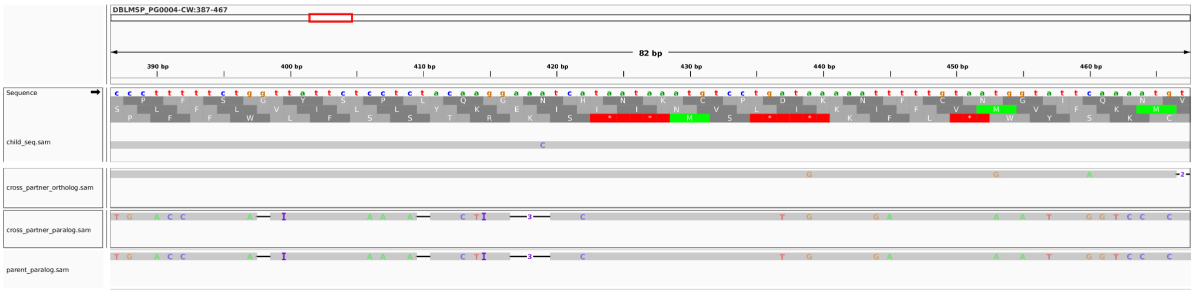


Supplementary Figure 5.1. **Two SNPs identified in one genetic cross progeny.** In one progeny sample from genetic cross HB3xDd2, two SNPs were identified in the DBs, one in each gene (panel a: DBLMSP2, panel b: DBLMSP). In both panels, the top track shows the parent gene sequence (HB3), and four subsequent tracks are shown below, each representing one different aligned sequence (grey horizontal bars). The first of these is the child sample gene sequence, showing a single SNP difference to the parent. To confirm these were spontaneous mutations and not single-base gene conversions from an ortholog, three orthologous sequences that could have been conversion donors were aligned to the parent: the orthologous sequence from the other cross parent (Dd2, second track), and the paralogous sequences from both parents (third and fourth tracks). No matches to these at the SNP positions can be seen.

### A new mechanism for allelic dimorphism

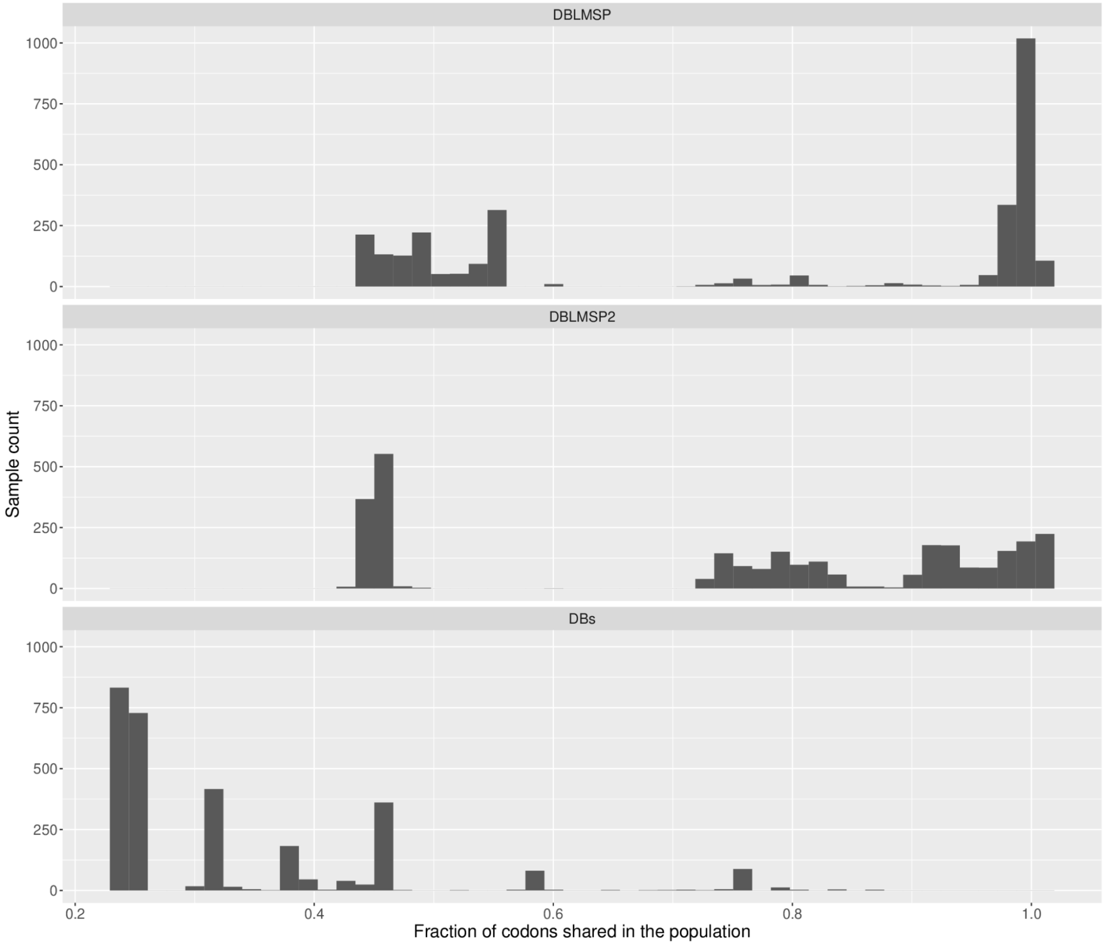


Supplementary Figure D.1. **Diversity and divergence levels in the DBL-spanning region (DSR) of the DBs**. The two first panels measure, for each of DBLMSP and DBLMSP2, the percent codon identity of 2,882 randomly-chosen gene pairs, and is a measure of sequence diversity. The third panel shows the percent codon identity between DBLMSP and DBLMSP2 across all 2,882 samples where they were confidently-resolved, and is a measure of sequence divergence. Between-gene divergence exceeds within-gene diversity (lower codon identity across genes than within genes).

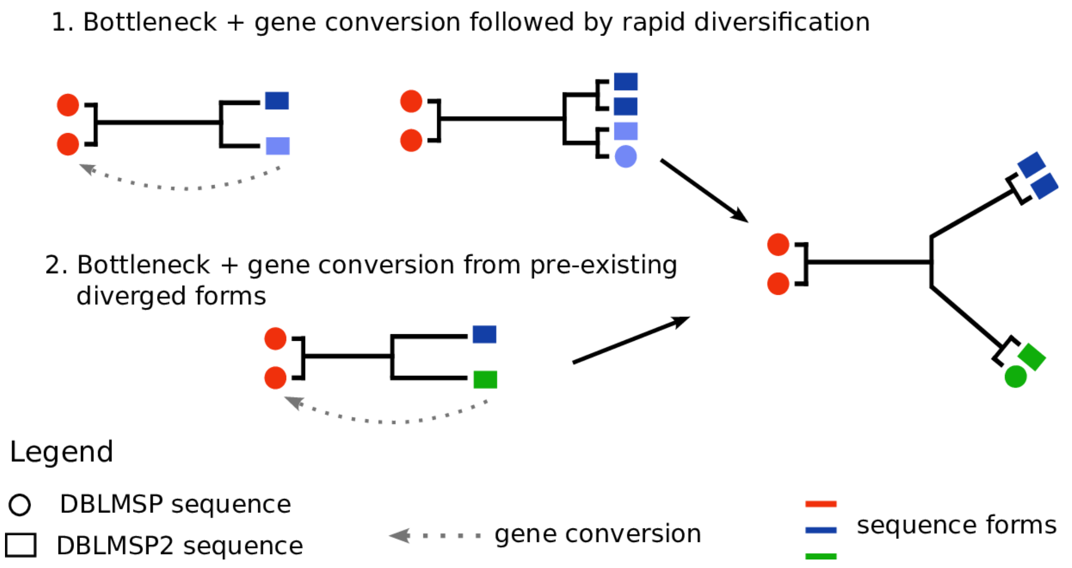


Supplementary Figure D.2. **Two evolutionary scenarios consistent with the observed data in Figure 6.** Both rely on two assumptions. First, that the DBs diverged from each other at a faster rate than either gene diversified (as expected under both diversifying selection or simple genetic drift), which we found applied (supp. fig. D.1). Second, that a population bottleneck occurred at some time in the past, as clearly supported by previous studies (16,17). Then we propose that following a strong bottleneck, gene conversion created allelic dimorphism in one of the two genes. Given that, in Figure 6, clade B is closer to clade A than clade C, we assume conversion occurred from DBLMSP2 to DBLMSP, creating dimorphism in DBLMSP. To explain the dimorphism in DBLMSP2, we propose that either DBLMSP2 rapidly diversified from a single or a few closely-related lineages (scenario ‘1’), or that two pre-existing, highly-diverged DBLMSP2 sequences were sampled during the bottleneck (scenario ‘2’). Low recombination rates between forms would be expected given low homology levels. Importantly, neither scenario requires that exactly two forms exist in each gene in the long-term nor that recombination does not occur between forms, and both explain dimorphism as a product of gene conversion between diverged paralogs following a bottleneck.
